# Not optimal, just noisy: the geometry of correlated variability leads to highly suboptimal sensory coding

**DOI:** 10.1101/2022.03.08.483488

**Authors:** Jesse A. Livezey, Pratik S. Sachdeva, Maximilian E. Dougherty, Mathew T. Summers, Kristofer E. Bouchard

## Abstract

The brain represents the world through the activity of neural populations. Correlated variability across simultaneously recorded neurons (noise correlations) has been observed across cortical areas and experimental paradigms. Many studies have shown that correlated variability improves stimulus coding compared to a null model with no correlations. However, such results do not shed light on whether neural populations’ correlated variability achieves *optimal* coding. Here, we assess optimality of noise correlations in diverse datasets by developing two novel null models each with a unique biological interpretation: a uniform correlations null model and a factor analysis null model. We show that across datasets, the correlated variability in neural populations leads to highly suboptimal coding performance according to these null models. We demonstrate that biological constraints prevent many subsets of the neural populations from achieving optimality according to these null models, and that subselecting based on biological criteria leaves coding performance suboptimal. Finally, we show that the optimal subpopulation is exponentially small as a function of neural dimensionality. Together, these results show that the geometry of correlated variability leads to highly suboptimal sensory coding.

## Introduction

The brain represents the world through the coordinated firing of neural populations. For instance, neural populations in early sensory areas are thought to transform the features of stimuli and transmit them to downstream cortical areas. Indeed, many studies of sensory areas seek to analyize what sensory features are transmitted in the brain and with what fidelity. Understanding population neural activity necessitates analyzing the joint activity of many neural units, beyond single-neuron analysis. Normative theories, which formalize optimality criteria, are powerful tools in these analyses, as they can establish principles for explaining features of experimentally observed neural activity at the population level. Therefore, it is important to develop methods for quantitatively assessing normative theories based on the features observed in neural data. One prominent feature of neural activity is variability: neural recordings exhibit trial-to-trial fluctuations in response to the same stimulus. From a normative perspective, the geometry of variability in neural activity impacts how optimally a population of neurons can encode stimuli [1, 2]. However, the optimality of correlated variability has not been assessed.

Many studies have found pairwise correlations in the trial-to-trial variability of the firing rates of simultaneously recorded neurons, often called correlated variability or noise correlations [3–9]. The correlated variability observed in experimental studies typically depends on the tuning and stimuli [10–12]. For example, **Figure 1a, b** shows the single-trial variability in Ca^2+^ responses (∆F/F) for two simultaneously recorded mouse retinal ganglion cells (RGCs) in response to drifting bars. The RGCs’ correlated variability for a single stimulus is shown in **Figure 1e**. Although correlated variability is typically considered in simultaneous single neuron electrophysiology measurements, it has been observed in calcium imaging recordings [13] and larger scale measurements such as electrocorticography recordings [9]. Correlated variability has many possible biological sources in neural populations (see **Supplementary Fig. 1**) [6, 7, 14–18], which the nervous system may be able to modify. Understanding the impact of correlated variability on population coding is important for revealing the principles governing neural computation [1, 2, 4].

**Figure 1:**
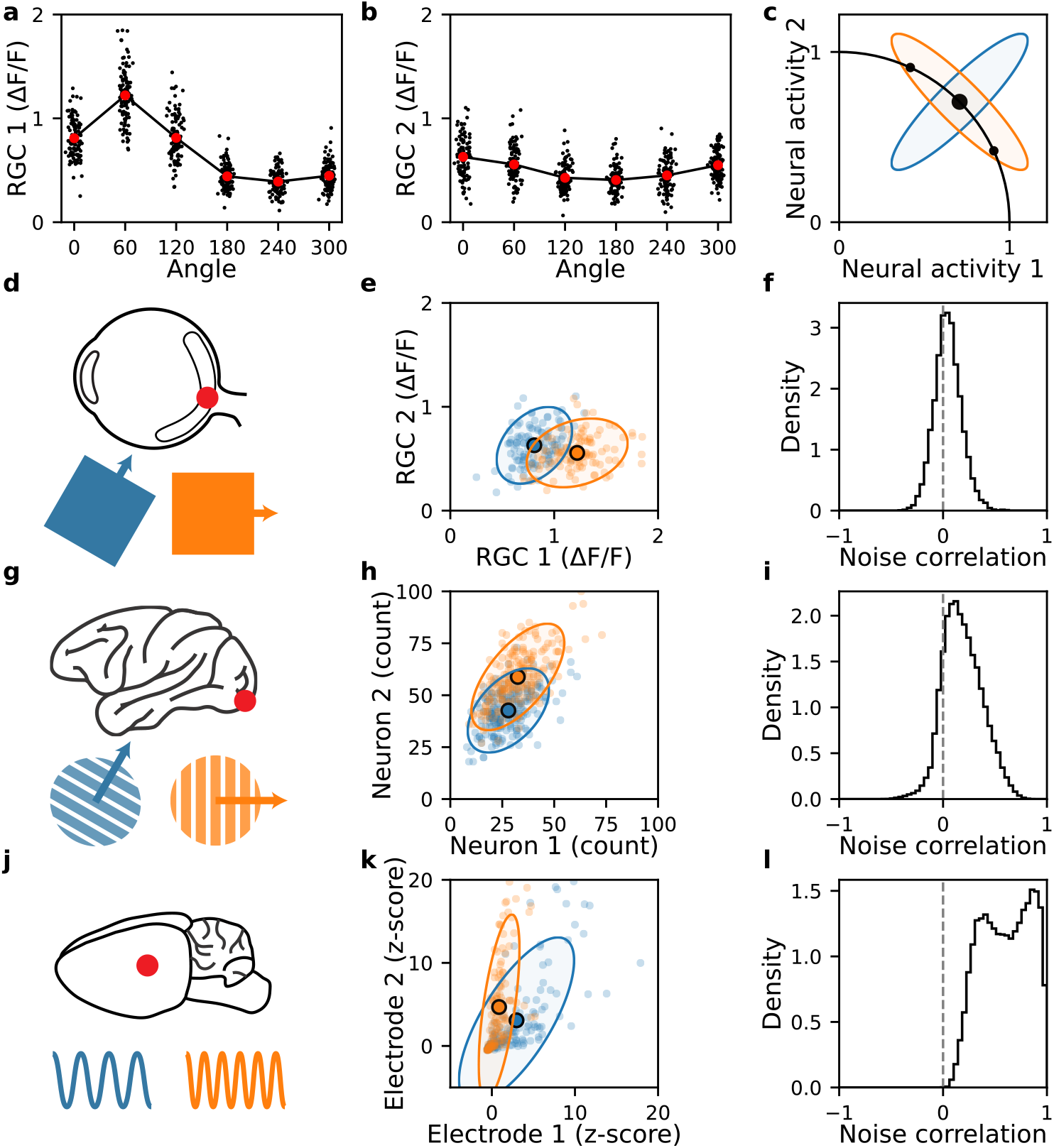
Correlated variability is a pervasive neural phenomenon. **a, b.** Mean activity as a function of bar angle (larger open circles) and trial-to-trial variability (small dots, small angle offsets for visualization) for angle 0 (corresponds to the blue dots for Neuron 1 and Neuron 2, respectively, in **d**). **c.** Illustration of mean stimulus response curve (black line), less detrimental correlated variability (blue ellipse), and more detrimental correlated variability (orange ellipse) for two model neurons. The large black dot is the mean stimulus response corresponding to the covariances. The small black dots are the mean responses for neighboring stimuli. **d-l.** Each row refers to a different experimental dataset, while columns refer to an aspect of the dataset. **d-f.** Calcium imaging recordings from mouse retinal ganglion cells in response to drifting bars. **g-i.** Single-unit spike counts recorded from primary visual cortex of macaque monkey in response to drifting gratings. **j-l.** Micro-electrocorticography recordings (*z*-scored H*γ* response) from rat primary auditory cortex in response to tone pips at varying frequencies. First column (**d, g, j**) depicts the recording region and stimulus for each dataset. Second column (**e, h, k**) shows the activity of two random RGCs/neurons/electrodes in the population to two neighboring stimuli. Individual points denote the unit activity on individual trials, while covariance ellipses denote the noise covariance ellipse at 2 standard deviations. Third column (**f, i, l**) plots the distribution of pairwise noise correlations, calculated for each pair of units across stimuli.

Correlated variability impacts the fidelity of a neural code when discriminating stimuli. Theoretical and computational studies have determined how the interplay between correlated variability and tuning properties affect population coding [2, 8, 15, 19–23]. **Figure 1c** shows the mean response curve (black line, defined by the mean firing rate of the neurons in response to various stimuli) from two hypothetical simultaneously recorded neurons across a range of stimulus values (3 neighboring stimuli are demarcated with black dots). From a geometric perspective, if the correlated variability has low variance (**Fig. 1c**, blue ellipse) along the mean stimulus response curve (**Fig. 1c**, black line), the impact on coding will be less detrimental than having high variance (**Fig. 1c**, orange ellipse) along the stimulus response curve. This is because the trial-by-trial fluctuation (blue ellipse) in response to the central stimulus (large black dot) will minimally overlap with the response to the nearby stimuli (small black dots). In early sensory areas, such as retina and primary visual cortex, studies have found that correlated variability enhances population coding [6, 16, 24–28]. Outside of early sensory areas, both the structure of correlated variability and its impact on coding is heterogeneous [12, 29, 30]. Brain states can change correlated variability and therefore its effect on population coding [31–33]. These studies leave open the possibility that the correlated variability is optimal for coding in sensory areas, which has not been evaluated.

The impact of correlated variability on neural coding is typically assessed by comparing the linear Fisher information (LFI) of the experimentally observed correlations to the distribution of LFI under the *shuffle null model*, a null distribution with the same per-neuron variability, but no correlations across neurons. LFI quantifies how accurately neural population activity can be used to distinguish two stimuli. Many previous studies have shown, by using the shuffle null model, that the geometry of correlated variability can benefit neural coding. However, comparing the experimentally observed correlated variability with the zero correlation version is only one relevant comparison for determining optimality; there are potentially other geometries which are not captured by the shuffle null model. In principle the brain’s correlated variability could have produced better (or worse) coding properties. Furthermore, it is unclear whether zero-correlation population activity is the only reasonable null distribution given biological processes such as learning, highlighting the importance of developing tailored null models [34]. Testing normative theories of stimulus coding in neural datasets requires understanding whether the geometry of experimental correlated variability is optimal, however methods for testing the optimality of correlated variability are currently lacking.

In order to test the optimality of correlated variability in experimentally observed neural responses, we developed two null models. The *uniform correlation null model* and the *factor analysis null model* each define a null distribution of correlated variability and have a particular biological interpretation. Using these null models, we test the optimality of neural coding in newly acquired data recorded from retinal ganglion cells (RGCs, Retina), previously recorded neurons in primary visual cortex (V1), and newly acquired ECoG electrodes on primary auditory cortex (PAC) (**Fig. 1d-l**). These datasets span neural areas and recording modalities used in many previous studies. Our main finding is that the experimentally observed geometry of correlated variability leads to highly suboptimal coding across all datasets and both null models. Furthermore, the degree of suboptimality worsens as a function of the number of neural units considered in the neural population. We find that for a large fraction of subsamples of the recorded units, achieving optimality would push the neural responses into regimes that violate biological constraints. However, even when neural units are subsampled to optimize for biological criteria, they remain highly suboptimal. Finally, direct selection of optimal subsamples shows that the optimal population is exponentially small as a function of neural dimensionality. Our results demonstrate that the traditional null model of correlated variability cannot be used to assess the optimality of neural data, and that biological constraints limit the ability of neural activity to achieve optimal correlated variability as defined by our null models. Together, our results show that the geometry of correlated neural variability leads to highly suboptimal sensory coding.

## Results

In order to assess the optimality of correlated variability in neural populations, we used three neural datasets which span animal models, sensory recordings areas, and recording modalities (**Fig. 1**). The newly recorded retina dataset is calcium imaging recordings in mouse retinal ganglion cells (RGCs) (**Fig. 1d-f**). The stimuli are drifting bars at 6 angles with each stimuli being presented 114 times. The previously recorded V1 dataset is spike sorted, single unit electrophysiology recordings in macaque V1 (**Fig. 1g-i**) [35]. The stimuli are drifting gratings at 12 angles with each stimuli being presented 200 times. The newly recorded primary auditory cortex (PAC) dataset is high gamma amplitude from *μ*ECoG recordings in rat primary auditory cortex (**Fig. 1j-l**). The stimuli are tone pips at 30 different frequecies with each stimuli being presented 60 times. We will refer to RGCs/neurons/electrodes as neural units. The neural units have various levels of pairwise noise correlations, *ρ*, across datasets (**Fig. 1f, i, l**), which is a key quantity for analyzing correlated variability. See Methods for more details on dataset recording and preprocessing.

### Methods for assessing the optimality of neural codes

An abundance of work has aimed to assess whether observed correlated variability is beneficial or detrimental for neural coding[6, 9–12, 16, 19, 24–33]. These studies often quantify the discriminability or fidelity of a neural code with the linear Fisher information (LFI, see Section) [36], which is a measure of how well the neural activity could be used to discriminate between different stimuli. The LFI is a function of the stimulus, *s*, the stimulus-derivative of the mean neural activity, 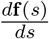, and the variability of the neural activity tahroeumnedan, Σ(*s*), and can be written as:

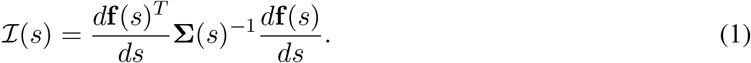

Typically, the impact of correlated variability is assessed by comparing the experimentally observed LFI to a distribution of LFIs generated from the shuffle null model. Trial-shuffling the data will produce a distribution over covariance matrices (Σ(*s*)) where the pairwise correlations are all centered near zero (**Fig. 2a**, observed covariance is filled, corresponding shuffle covariance is dashed). However, the shuffle null model does not compare the observed correlations to a broad range of potential non-zero correlations. In principle, neural circuits can support a range of covariance structures with significant nonzero pairwise correlations, many of which can produce higher LFI than having zero correlations. In this case, using the shuffle null model would overestimate the level of optimality in neural data, and therefore cannot be used to assess the optimality of the experimentally observed correlations. To our knowledge, the optimality of correlated variability has not been evaluated on neural data before.

**Figure 2:**
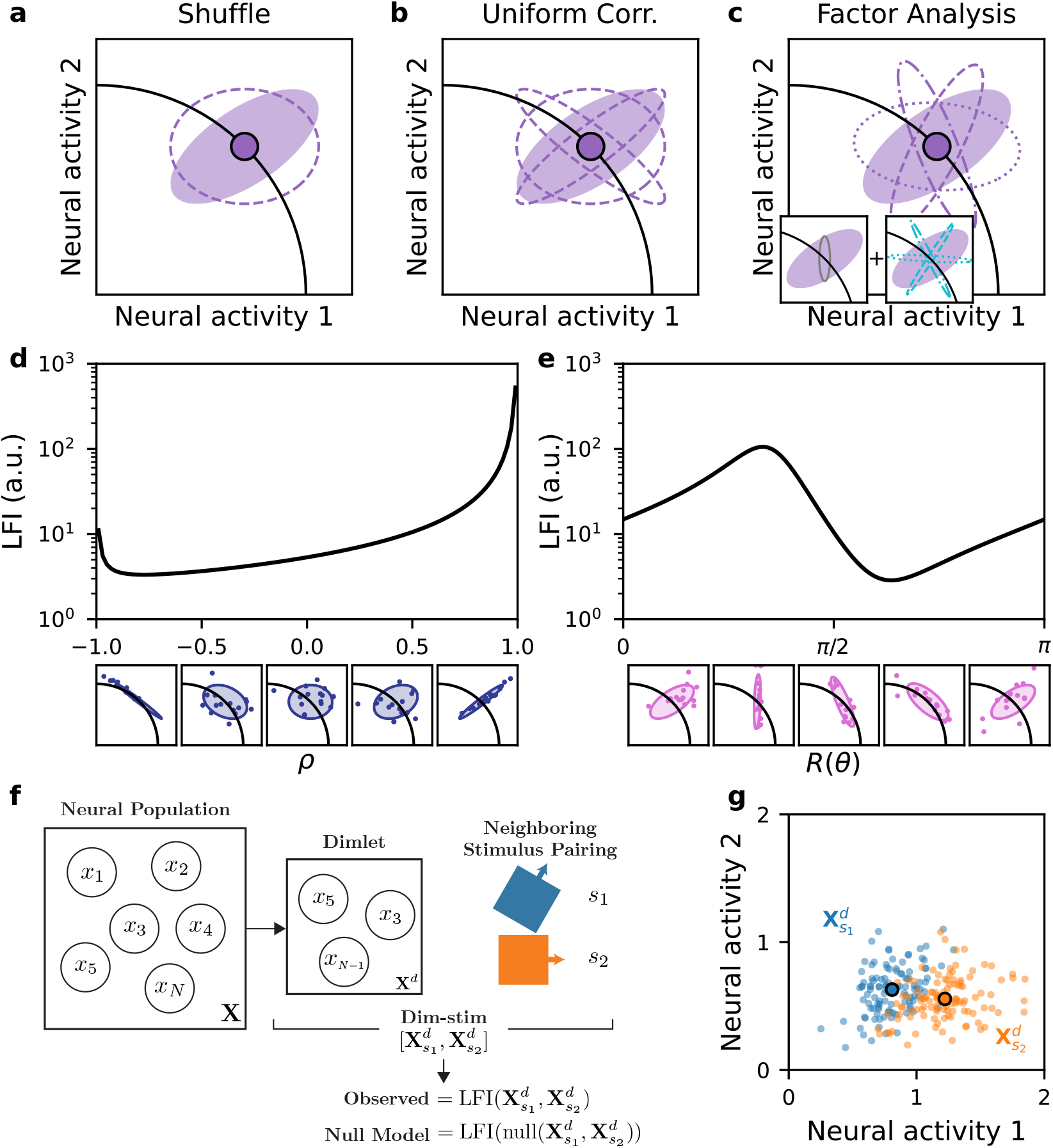
Methods for assessing the optimality of neural codes. **a-c.** Null models of correlated variability. Solid, purple ellipses denote the trial-to-trial variability observed about the mean stimulus activity (solid point). Samples from the null models are depicted by dashed ellipses. **a.** The shuffle null model maintains per-neuron variance and samples correlations near zero. **b.** The uniform correlation null model maintains per-neuron variance and samples uniform correlations. **c.** The factor analysis null model combines a fixed private variability (estimated from the experimental data, left gray inset) with shared variability (right teal inset) that can be rotated to form null samples (dash styles are consistent between the teal shared variabilities in the inset and the purple null samples in the main panel). **d.** For a synthetic 2d dataset, the LFI for the fixed-marginal parameterization as a function of the pairwise correlation, *ρ*, is shown at the top, the bottom plots are the covariance and samples as a function of *ρ*. **e.** For a synthetic 2d dataset, the LFI for the factor analysis parameterization as a function of the rotation angle, *θ*, is shown at the top, the bottom plots are the covariance and samples as a function of *θ*. **f.** To calculate an observed LFI or percentile under a null model, *d* units were randomly drawn from the population to form a “dimlet”. Then, two neighboring stimuli, *s*_1_ and *s*_2_, were chosen. The dimlet and stimulus pairing together constitute a “dim-stim”, or a pair of design matrices 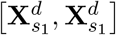. These dim-stims are the samples inputs into a LFI calculation or null model analysis and form the basis for distributions of calculated quantities. **g.** Dim-stim responses in the retinal data for the depicted stimulus pairing (colors) from **f.**

In order to assess optimality, the null model should be chosen to adequately span achievable covariance structures. Defining achievable may depend on the experimental context, including the types of neurons being recorded, their location in the brain, or the recording modality. Thus, it is beneficial if the parameters of the null model have a biological interpretation. We propose two null models that allow us to asses the optimality of experimental neural responses: the uniform correlation (UC) null model and the factor analysis (FA) null model. The uniform correlation null model maintains the per-neural unit distributions of activity, like the shuffle null model. In contrast to the shuffle null model which samples the correlations around zero (**Fig. 2a**), the uniform correlation null model samples the multivariate correlations uniformly (**Fig. 2b**, dashed lines are samples with different correlations) [37]. In the UC null model, neural units maintain their private mean and variance for a particular stimulus, but have the freedom to change their multivariate pair-wise correlations (*ρ*). Biologically, changing the pairwise correlations could be achieved through recurrent connectivity within the network of neural units. Depending on the correlations, the network could achieve a range of coding fidelities as assesed by the LFI (**Fig. 2d**, covariance structures shown below the plot lead the LFI as a function of the scalar pairwise correlation (*ρ*). At extreme values of correlation, the LFI can take on the highest values [22]. Mathematically, the UC null model constrains the per-unit variances while sampling the multivariate correlations (*ρ*) uniformly (see Methods for details). Motivated by experimental findings that the variability in population responses has private and shared components [7, 14], we also developed a factor analysis (FA) null model. The FA null model decomposes the experimentally observed covariance into independent private variances and shared variability [7, 15]. The private variance is fixed (**Fig. 2c**, gray ellipse in the left inset) and the shared variability’s weighting on different neural units can change through a rotation (**Fig. 2c**, dashed teal ellipses in the right inset are sampled rotations of the shared variability). Biologically, this models each neuron having fixed private variability and incoming shared variability which could be weighted in different ways. As the shared variability is rotated, the covariance structure varies, and the LFI takes on a smaller range of values than in the UC null model (**Fig. 2e**, covariance structures shown below the plot generate the LFI as a function of rotation angle, *R*(*θ*)). Mathematically, the FA null model constrains the factor analysis private variances but applies uniformly sampled rotations to the loading matrix for the shared variability (see Methods for details). Together, these null models define the potential space of covariances based on two different biological motivations and provide suitable tests of optimality.

To use the null models, for each neural population (retinal ganglion cells, V1 neurons, electrodes in primary auditory cortex), we randomly sampled “dimlets”, or sub-populations of neural units, of dimension *d*. We combined dimlets with a variety of neighboring stimulus pairings to obtain a subset of the neural responses which we call a dim-stim (**Fig. 2f**, see Methods). A dim-stim would be the input to the task of constructing a decoder for neighboring stimuli using a neural sub-population’s responses across trials (**Fig. 1e, h, k** and **Fig. 2g**). For each dataset, we generated a large number of dim-stims across a set of dimensions *d* = 3 – 20 (see Methods). We calculated the LFI for each dim-stim across dimensions and datasets. We refer to this quantity as the observed LFI. Next, we sampled the null models 1,000 times for each dim-stim, and calculated the LFI for each sample (see Methods). Thus, for each dim-stim, we obtained a single experimental LFI and a corresponding distribution of LFIs for each null model. The 1,000 null LFIs constitute a null distribution to compare the experimentally observed LFI against. In particular, we define the percentile as the fraction of the 1,000 null LFIs which are less or equal to than the observed LFI. Higher percentiles indicate that the observed LFI is larger than more samples from the null model.

### The geometry of correlated variability leads to suboptimal neural coding

With the uniform correlation (UC) and factor analysis (FA) null models, we assessed the optimality of the neural code. To characterize the optimality of a wide range of sub-population and stimulus settings, we performed a large scale analysis evaluating the LFI in both the experimentally observed data and null models (see Methods). We compared the experimentally observed LFI to the distribution of LFI from the null models. Specifically, for the experimental data, we compute the median LFI across dim-stims at each dimension (**Fig. 3a-c**, black lines). For the shuffle, uniform correlation (UC), and factor analysis (FA) null models, we first calculated the median LFI from the null distribution for each dim-stim and then report the median across dim-stims (**Fig. 3a-c**, gray, blue, and orchid lines, respectively).

**Figure 3:**
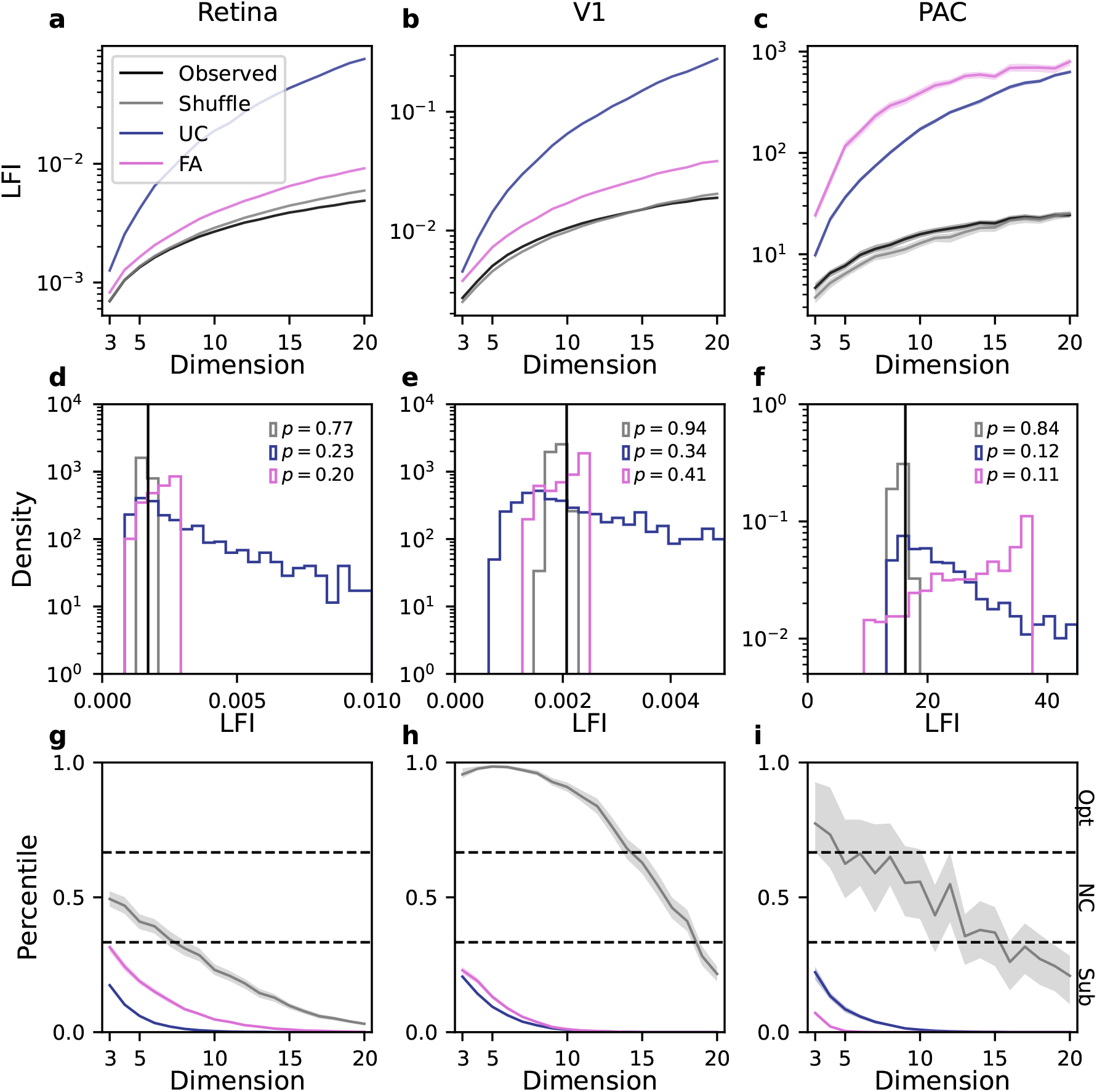
The geometry of correlated variability leads to suboptimal neural coding. Each column corresponds to one of the datasets. Color legend is shared across columns. Color legend is preserved across panels. **a-c.** The median LFI is plotted (solid lines, log-scale *y*-axis) as a function of the dimlet dimension (*x*-axis) for the observed correlated variability and null model samples (colors in legend). Shaded regions indicate the 95% CI of the median LFI (note that CIs are often comparable to the median line width). **d-f.** Histograms of null LFIs are shown for the shuffle, uniform correlation, and factor analysis null models for one dim-stims for each dataset. The observed LFI is denoted by the black vertical line in each plot. Percentiles for each null model are reported. **g-i**. Median observed dim-stim percentiles are shown (solid lines) as a function of dimlet dimensions, for each dataset and null model. Shaded regions indicate the 95% CI of the median observed percentile (note that CIs are often comparable to the median line width). Black dashed lines divide optimal (Opt), near-chance (NC), and suboptimal (Sub) regions.

As expected, the experimentally observed LFIs across dim-stims grew with dimlet dimension, indicating that increasing the dimension of the neural population improved the stimulus decoding (**Fig. 3a-c**, black lines). Similarly, the median null model LFIs grew with dimlet dimension. The shuffle null model exhibited comparable discriminability relative to the experimental LFI at lower dimensions (**Fig. 3a-c**, gray lines). At higher dimensions, however, the shuffle null model LFIs began to exceed the observed LFIs. In contrast, both the uniform correlation and factor analysis null models exhibited considerably larger median LFIs than the observed data, with the disparity increasing with dimlet dimension. Therefore, on average, the stimuli were more easily discriminable using the covariances sampled from the UC and FA null models than the experimental covariance. We further observed differences across datasets. For example, the factor analysis null model (**Fig. 3a-c**, orchid lines) exhibited similar LFIs to the uniform correlation null model for the PAC dataset. However, in the retina and V1 data, the factor analysis LFIs were more comparable to the observed and shuffle LFIs. Overall, **Figure 3a-c** demonstrates that the uniform correlation and factor analysis null models produce LFIs that generally exceed the LFIs of the observed data, suggesting the neural code is suboptimal.

Although the differences between the null model LFIs and observed LFIs were large, the preceding analysis was done at a population level rather than comparing each dim-stim LFI with its own null distribution. Therefore, we quantified the optimality per dim-stim, relative to a null model, with its observed percentile. To calculate the population optimality measure, the median percentile across dim-stims is taken. A higher percentile means that the observed LFIs are greater than a larger fraction of the null LFIs. To operationalize the notion of population optimality, we define three categories for optimality based on the median of the experimental distribution of percentiles. If the median is greater than 2/3, the population is optimal (Opt), if the median is between 1/3 and 2/3 the population is near-chance (NC), and if the median is below 1/3 the population is suboptimal (Sub). Alternative categorizations could be used, but we chose the even splitting into thirds for simplicity (see Methods for details).

We found that each null model exhibits distinct LFI distributions, with further variation depending on the dataset and dim-stim. Example null model distributions for individual *d* = 3 dim-stims are depicted in **Figure 3d-f** (vertical black line indicates the experimental LFI, gray, blue, orchid are the shuffle, UC, and FA null model LFI distributions respectively, note that the uniform correlation null distributions often have long tails and are truncated for visualization). The examples highlight that the percentiles can vary across null models for a dataset (**Fig. 3d-f**, inset text). The heterogeneity in observed percentiles motivated examining their distribution across all dim-stims. Thus, for each dataset, we computed the distribution of observed percentiles across the dim-stims per dimlet dimension (*d* = 3 to *d* = 20). The median observed percentile (calculated across dim-stims) as a function of dimlet dimension is shown in **Figure 3g-i**. Consistent with other studies [6, 13, 16], we found that the shuffle null model (gray lines) often had large observed percentiles, indicating that the shuffle null model often showed the benefits of experimentally observed correlations versus having no correlations. However, it would be misleading to interpret these results as a test of optimality. Indeed, compared to the uniform correlation (blue lines) and factor analysis (orchid lines) null models, the experimental data exhibited suboptimal observed percentiles (**Fig. 3g-i**, blue and orchid lines). All percentiles decreased with dimlet dimension, implying that the neural representations became less optimal as the number of neurons increases. In theory, this decrease is expected as eventually differential correlations induce information saturation in the populations, however recent work indicates that we should not expect to see the impact of differential correlations at this relatively small scale [38–40]. Indeed, saturation of the LFI was not evident in **Figure 3a-c**. This indicates that the suboptimality observed in **Figure 3g-i** is not due to differential correlation, but from some other biological cause.

**Figure 3g-i** also highlights differences across datasets. The shuffle null model had the lowest observed percentiles among the three datasets for the retina data, starting near-chance for small dimlet sizes and dropping below 1/3 around *d* = 7 (**Fig. 3g**, grey lines). For the V1 data, the shuffle null model clearly exhibited the highest observed percentiles, indicating the coding benefits of correlations compared to zero correlations for small dimlet sizes up to *d* = 15 (**Fig. 3h**, grey lines). In the primary auditory cortex data, the shuffle null model exhibited intermediate observed percentiles, with a larger spread in confidence intervals, indicating a higher heterogeneity in the observed percentiles (Fig. 3i, gray shaded region). Meanwhile, the observed percentiles for the uniform correlation and factor analysis null models were more similar across the three datasets, with slightly different magnitudes. In particular, the retinal data exhibited the largest observed percentiles for the factor analysis null model, while the PAC data exhibited the smallest, going to zero around *d* = 5. The uniform correlation null model had the lowest percentiles for the retina dataset and similar percentiles for the V1 and PAC datasets. This behavior roughly tracked the distribution of pairwise correlations amongst the three datasets (**Fig. 1e, h, k**), with the retinal data possessing the lowest average noise correlation, and the PAC data possessing the highest average noise correlation. Critically, across all datasets and dimensions, the percentiles for both the uniform correlation and factor analysis null models were below 1/3. This indicates that the geometry of correlated variability leads to suboptimal coding, and that the suboptimality becomes more pronounced with increasing neural dimension.

### Optimal correlated variability is typically biologically inaccessible

The results of the preceding section indicate that the geometry of correlated variability is highly suboptimal, as opposed to near-chance or optimal. We next sought to understand why this was the case. For the uniform correlation model, we summarize findings about optimal correlations from Hu *et al.* [22]. For the factor analysis model, we compared the structure of the observed covariances to those of the optimal covariances.

When the per-neural unit variability is fixed, as in the shuffle and uniform correlation null models, Hu *et al.* [22] showed that the optimal covariance structure will lie on the boundaries of the allowed values of *ρ* for several measures of coding fidelity, including the LFI (**Fig. 2d**). The authors discussed that points on the boundary may fall outside of biologically allowed regions. Consistent with this, we found that optimal correlation matrices for the uniform correlation null model often had absolute pairwise correlations that are close to 1, which was never observed in the experimental data (see **Supplemental Fig. 3**). Thus, the optimal correlated variability structure suggested by the uniform correlation null model may be biologically inaccessible. Meanwhile, the factor analysis model allows the distribution of highest pairwise correlations to be modified (and generally increased), but does not extend near 1, suggesting that the distribution of noise correlations achieved by the factor analysis null model is more biologically realistic.

Both the shuffle and uniform correlation null models will necessarily reproduce the observed single-unit statistics, because they only change the correlations. Therefore, both of these null models will reproduce the Fano factors 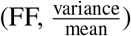 and negative densities (ND, fraction of activity below the smallest responses of the experimental activity) of the observed data. The factor analysis null model, however, can produce covariance ellipses that have different single-unit distributions. Thus, some FA-optimal covariances may orient variance in the negative or low-activity regions of the neural space. For the factor analysis null model, we quantified the degree to which the biological inaccessability of optimal covariances related to the percentiles of the experimental data for each dim-stim. The Fano factor quantifies the variability of neural units relative to their average activity. Typically, Fano factors for single-unit firing rates have been observed to be near 1 [41–44], in line with the approximately Poisson nature of firing rates. Thus, a large deviation from the Fano factors observed in the experimental data indicates the single-unit properties of the optimal covariances are biologically implausible (**Supplementary Fig. 2**). First, we examined whether the observed Fano factor diverged from the Fano factors achieved by the FA-optimal covariance on each dim-stim via their absolute log-ratio (see Methods). Large values of this quantity indicate greater difference between optimal and experimental single-unit distributions, suggesting less biological plausibility. Relatedly, a sample-covariance that has negative neural activity can be interpreted as less biologically plausible, because negative activity is either unachievable (for single-unit count variables) or highly unlikely (calcium imaging ∆*F/F* or baseline *z*-scored *μ*ECoG) (**Supplementary Fig. 2**). Therefore, the second quantity we examined was the absolute difference in negative density (ND), which captures the degree to which the FA-optimal covariance has negative neural activity (see Methods). Larger values of the negative density imply less biological plausibility. We used these two measures of biological plausibility to assess when the observed neural responses can be optimal according to the FA null model.

We determined whether the Fano factor (FF) and negative density (ND) distributions of the optimal covariances from the FA null model related to the suboptimality of the experimentally observed neural code. To do this, we directly compared the optimal FA null model Fano factors to the experimental Fano factors in **Figure 4a-c**. Across dim-stims, for *d* = 3, **Figure 4a-c** shows 2d-histograms of the absolute log-ratio of Fano factors against the FA percentile, with darker colors corresponding to higher log-density of samples. For each histogram, we additionally plot the median percentile as a function of the log-ratio in blue. We found that when the Fano factors closely matched (i.e., the log-ratio was close to zero), the percentiles spanned a broad range between 0 and 1 (medians percentiles: 0.51, 0.41, 0.15 for the lowest bin across datasets). However, FA-optimal covariances commonly deviated from the observed Fano factors, and when they did, the observed percentiles dropped below 0.5 and were often near 0. Thus, as the biological accessibility of the optimal covariance decreased, so did the optimality of the observed neural code. Likewise, for negative density (ND), we directly compared the optimal FA null model NDs to the experimentally observed NDs in **Figure 4d-f**. Across dim-stims, for *d* = 3, **Figure 4d-f** shows 2d-histograms of the absolute difference in NDs against the FA percentile with darker colors corresponding to higher log-density. For each histogram, we additionally plot the median percentile as a function of ND difference in red. We found that when the difference was close to zero, the percentiles spanned a broad range between 0 and 1 (medians percentiles: 0.47, 0.60, 0.31 for the lowest bin across datasets). However, the ND of FA-optimal covariances commonly deviated from the observed ND, and when they did, the experimentally observed percentiles were typically closer to 0.

**Figure 4:**
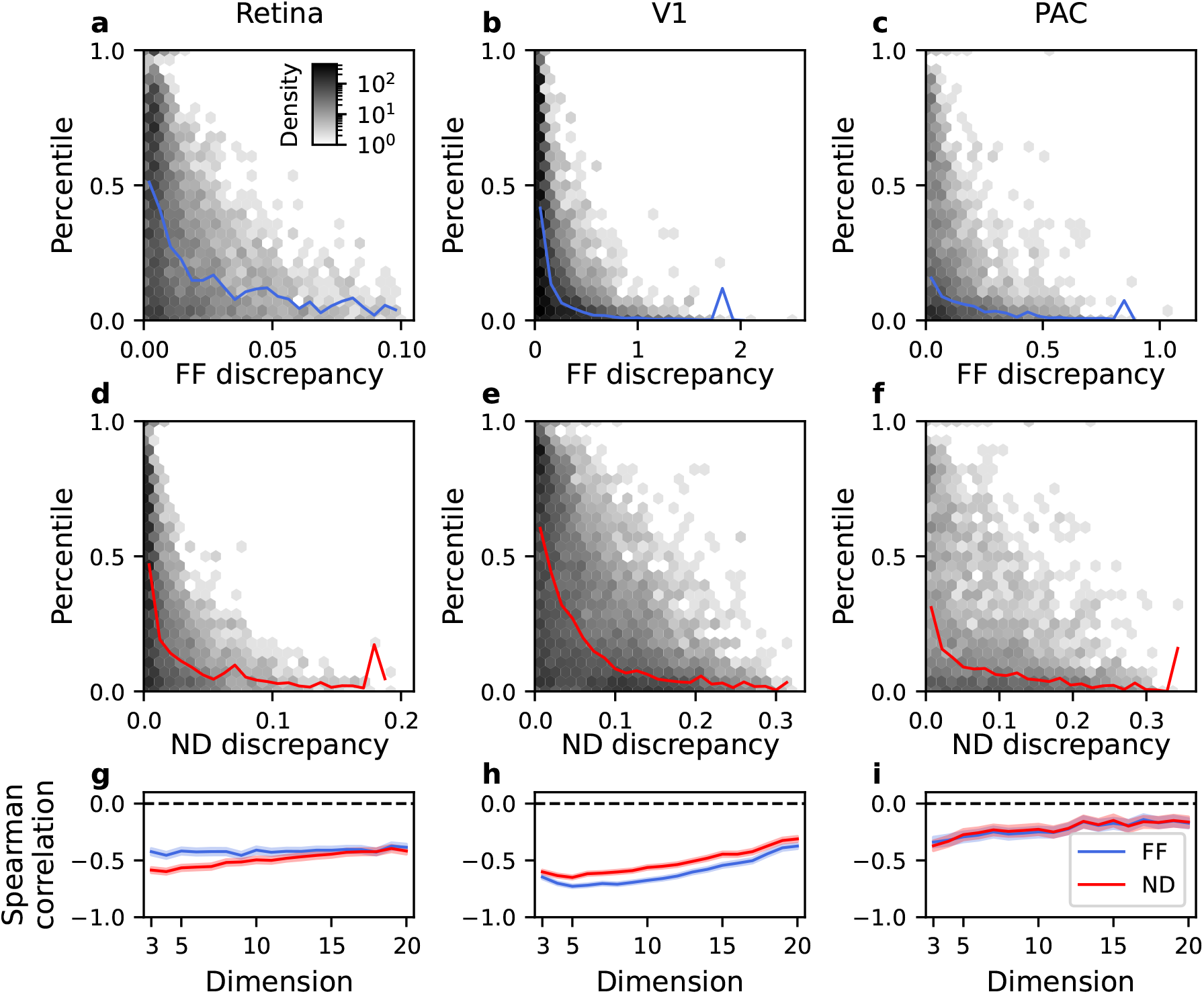
Optimal correlated variability is typically biologically inaccessible. Each column corresponds to a separate dataset. 2d histograms are plotted with a log-density color scale with shared colorbar. Color legend in **i** is shared across panels. **a-c.** 2d-histogram across dim-stims of the observed percentile under the FA null model versus the absolute log-ratio of the observed and FA-optimal covariance Fano factors for *d* = 3. Blue line is the median binned percentile as a function of the absolute log-ratio of observed and FA-optimal covariance Fano factors. **d-f.** 2d-histogram across dim-stims of the percentile under the FA null model versus the absolute difference of negative densities (ND) of the observed and FA-optimal covariance Fano factors for *d* = 3. Red line is the median binned percentile as a function of the absolute difference in NDs. **g-i.** The Spearman correlation coefficient between the observed percentile and absolute log-FF ratio or absolute difference of NDs, respectively is shown as a function of dimlet dimension. Dashed black line indicates zero correlation.

We summarized the relationship between biological plausibility and percentile for both FF and ND. At each dimension *d*, we calculated the Spearman rank correlation between the observed percentile and each measure of biological plausibility (**Fig. 4g-i**). For each dataset, we observed negative correlations that were significantly lower than zero across dimensions (*p* < 10^−5^, one sample *t*-test). These negative correlations imply that observed percentiles are smaller (i.e., the neural code is more suboptimal) when optimal correlated variability is biologically inaccessible. Together, these results indicate that the optimal covariances under the FA null model for *d* ≥ 3 are not biologically accessible.

### Optimal subpopulations are exponentially small

The results in the preceding section show that a majority of experimental dim-stims could not attain optimal covariances according to the UC and FA null models due to biological constraints. However, it is possible that although a majority of experimental dim-stims are suboptimal, there is a subset that are optimal, and these specific subpopulations are somehow utilized by the nervous system. If this was the case, the uniform sampling strategy over neural units may underestimate optimality as utilized by the nervous system. For example, in the retina, if we are imagining a downstream region like V1 is decoding the stimuli, then a more retinatopic sampling strategy, where retinal ganglion cells are more likely to be considered in a dimlet if they are located spatially near each other in the retina would be preferable. Alternatively, synaptic learning rules in downstream areas may select for neural populations that are tuned for similar stimuli. The responses to the preferred stimuli would be high and therefore we expect less Fano factor and negative density violation. Thus, it is possible that dim-stims subselected by these criteria will be more optimal than dim-stims sampled uniformly.

To test if biologically motivated subsampling of dim-stims improved the percentiles, we performed distance- and tuning-based subselection of the neural populations. For the retina and PAC datasets, we had access to the spatial locations of the RGCs/electrodes. We subselected 10% of dim-stims with the smallest average physical distance. Similarly, we subselected the 10% of dim-stims that had the most preferred stimuli (see Methods for details on subselection). We found that distance-based subselection did not reveal an optimal or near-chance subset of dim-stims (**Fig. 5a, c**, dotted lines and hatched shaded regions). Similarly, for the retina and V1 datasets, the tuning-based subselection did not reveal an optimal subset of dim-stims and the percentiles only improved to near-chance for the PAC dataset at *d* = 3 (**Fig. 5a-c**, solid lines and shaded regions). Furthermore, subselection directly based on the FF and ND criteria also did not find optimal or near-chance percentiles (**Supplementary Fig. 4**).

**Figure 5:**
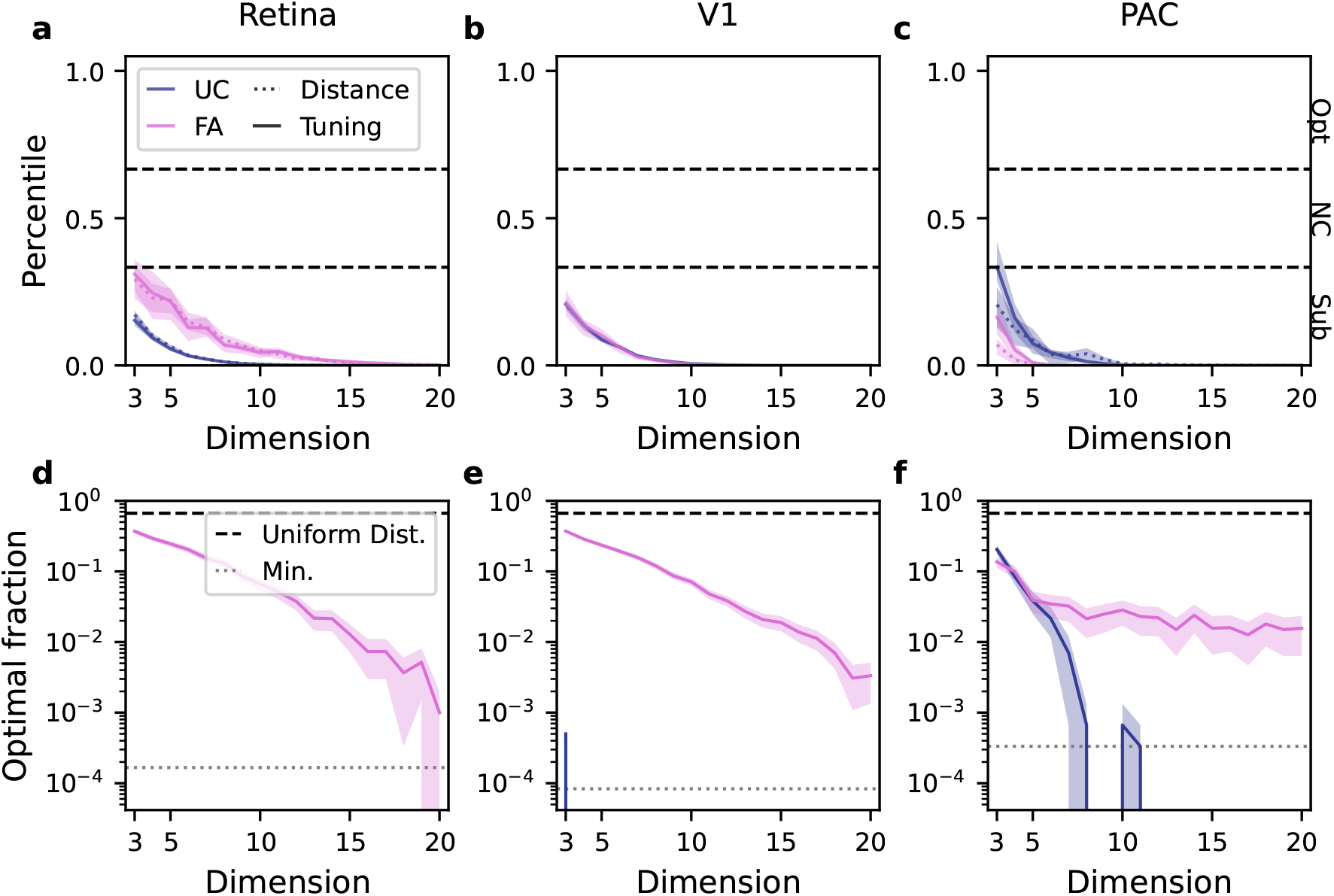
Optimal subpopulations are exponentially small. Color legend in **a** is shared across panels. a-c. For the uniform correlation and FA null model, dim-stims were subselected to maximize the units’ tuning (solid lines, highest 10% subselected). Additionally, for the retina and PAC datasets, dim-stims were subselected to minimize the average pairwise distance between the RGC RoIs in a dim-stim (dashed lines, lowest 10% subselected). The median percentiles are shown as a function of dimension. Black dashed lines indicate the 1/3 and 2/3 percentile range. Shaded regions indicate the 95% CI of the median percentiles. **d-f.** For each dimension, the largest possible fraction of dim-stim percentiles such that their median is 2/3 is plotted. Shaded regions indicate 95% CI. For the uniform correlation null model, dimensions where no samples exceeded the 2/3 threshold are not plotted. Black dashed line indicates the optimal fraction if percentiles were drawn from a uniform distribution. Gray dotted line indicates the minimum non-zero optimal fraction that can be estimated due to finite sampling.

Although these subselection criteria are biologically motivated, the previous results do not address whether any subpopulation of the neural units across stimuli have optimal percentiles, and if so, how small the subpopulation is. Intuitively, given the combination of a large enough neural population, variety of stimuli, and enough dim-stims, one would expect at least a small fraction of the dim-stims to have optimal percentile statistics by chance. To estimate the size of the optimal subpopulation, we calculated the optimal fraction of the neural population, that is, largest fraction of dim-stims that could be retained and still achieve optimal percentile statistics (median ≥2/3) (**Fig. 5d-f**). If the optimal fraction is smaller, optimal subpopulations are more rare. As a reference, if the distribution of percentiles was uniform, the largest two-thirds of the percentiles could be retained and their median would be 2/3, which is optimal (**Fig. 5d-f**, black dashed line). At *d* = 3 for the FA null model (**Fig. 5d-f**, orchid line), across datasets between 14% and 37% of the entire population was optimal if subselected. The optimal fraction according to the FA null model dropped below 10% by *d* = 4 − 9 and below 2% by *d* = 13 − 15 across datasets. At higher dimensions, the optimal subpopulation continued to become exponentially small, although the PAC dataset had a slower decrease. According to the uniform correlation null model, for the retina and V1 datasets, less than approximately 0.1% of the population was optimal since almost no subpopulation was found from the finite samples. At *d* = 3 for the PAC dataset, 20% of dim-stims would be considered optimal, but that drops below 1% by *d* = 7 and continued to decrease to the smallest possible estimated value by *d* = 12 since no subpopulations were found for higher dimensions. Finally, an alternative analysis of peaks in the percentiles near 1 in excess of what would be expected from a uniform distribution confirmed that there were exponentially small optimal populations (**Supplemental Fig. 5**). Together these results show that correlated variability is suboptimal in the neural recordings considered here. Furthermore, biologically motivated selection criteria are not able to find the exponentially small optimal subpopulations.

## Discussion

Determining the principles of the neural code is critical for a complete understanding of brain function. Correlated variability is prevalent in neural recordings and has been the subject of numerous studies seeking to understand its mechanistic sources and implication for neural coding. Many previous studies have found that the experimentally observed correlations can be a benefit to neural coding compared to having zero correlations [6, 13, 16, 24, 25, 27]. This suggests that the correlated variability could in fact be optimal. However, the shuffle null model used in these studies is not able to assess optimality. To the best of our knowledge, the optimality of correlated variability in neural data has not previously been assessed.

Here, we developed two null models which allow the optimality of observed correlated variability to be directly assessed: the uniform-correlation (UC) and factor analysis (FA) null models. Using these null models, we found that the experimentally observed neural activity across three datasets was consistently suboptimal. As more neural units were included in the neural population, the neural populations became more suboptimal. In order to more fully understand the suboptimality, we evaluated the characteristics of the optimal covariance and found that a consistent picture emerges: for a majority of neural subpopulations, the optimal covariance is biologically inaccessible. We then used biologically motivated subselection criteria to assess whether there were subpopulations with optimal coding statistics. We found that subsampling using criteria based on the tuning of units or the spatial location of the units does not result in increased coding optimality. Finally, we showed that optimal subpopulations based on *post-hoc* selection became exponentially small as the dimensionality of the neural population increased. Thus, we conclude that in the early sensory areas studied here, the geometry of correlated variability leads to highly suboptimal neural coding.

We observed suboptimal coding performance as assessed by both the uniform correlation and factor analysis null models. However, the magnitude of the suboptimality, as measured by the observed percentiles, differed across null models and datasets. The observed percentiles for the uniform correlation null model had a small trend from low to high for the retinal data, the V1 data, and the PAC data, respectively. This trend tracks with the distribution of noise correlations in each dataset (**Fig. 1f, i, l**), with the the retina dataset exhibiting, on average, the smallest magnitude noise correlations, and the PAC datasets exhibiting the largest. The smaller range of noise correlations exhibited by the retina suggests that there may be stronger biological restrictions on its correlated variability compared to V1 and PAC. The observed percentiles for the factor analysis null model trend from just below near-chance to highly suboptimal from retina to PAC. Thus, the larger correlations and more suboptimal coding performance indicates that shared variability in V1 and PAC is more likely to interfere with sensory coding. The retina and V1 recording modalities (calcium imaging and single-unit electrophysiology, respectively) measure putative single-unit activity where correlated variability in the recordings corresponds to correlated single neuron activity. Understanding the optimality of the neural code with these two modalities directly addresses decoding as a normative theory in early sensory areas. On the other hand, the correlated variability in the *μ*ECoG recordings in PAC is likely due to a combination of the correlations between the neural populations under each electrode and local tissue conduction [45, 46]. Due to this, the optimality of the high gamma amplitude correlated variability recorded with *μ*ECoG is a coarse-grained signal that may not be read-out by any downstream cortical area, but is important for understanding whether limitations in the accuracy of clinical ECoG-based brain-computer interfaces in humans may be due to correlated variability in the input signals.

Many studies of correlated variability, including ours, consider the impact of correlated variability from a decoding perspective. However, other normative perspectives exist. In Bayesian models of sensory processing [47], correlated variability could correspond to sampling from a relevant (posterior) distribution. In this case, correlated variability would be informative for understanding the structure of uncertainty in sensory processing, rather than nuisance variability as in the decoding perspective. Likewise, neural systems likely have other important constraints or ethological goals. Making decisions or generating behavior based on sensory information may be optimized by different correlation structures versus a purely decoding framework [48]. For example, Valente *et al.* [48] find that single-trial responses in posterior parietal cortex which have higher noise correlations also have more correct choices, contrary to expectation. They model this finding with a read-out network that computes an additional nonlinear “consistency” value across the population in addition to the linear sensory information for use in decision making. Huang & Lisberger [49] show that correlated variability in middle temporal visual area could plausibly be the cause of variability in smooth-pursuit eye movements. Even within the normative decoding framework, correlated variability which facilitates decoding as assessed by the LFI may not be the same as the correlated variability which facilitates information propagation or learning in more realistic nonlinear, noisy networks [5, 48, 50]. In these contexts, our formalism for creating null models could be used to test the optimality of neural codes, although as the assumptions on linear decoding are relaxed, it may become difficult to make theoretical predictions that hold generally.

The null models we proposed both have parameterizations that are interpreted in a fully Gaussian model. Generalized linear models [51, 52] or correlated multivariate distributions with binary-spike or spike-count distributions [53–55] could potentially better model nonlinearities between the parameters of the model and the non-Gaussian neural responses, which can impact estimates of neural coding optimality. In order to assess optimality in these models when fit to data, a similar formalism for generating null models is needed, where certain parts of the parameterization are fixed and others are given a null distribution. However, the independent parameterization of the mean responses (tuning) and correlated variability is a unique feature of the multivariate Gaussian distribution. Therefore, new analytical results would be needed to directly study the impact of non-Gaussian correlated variability on neural coding. A broader set of null distributions could similarly be used in phenomenological models of correlated variability which combine tuning and various types of (correlated) noise [6, 21, 28, 56] or in mechanistic models, which attempt to simulate some aspects of the neural circuit which lead to correlated variability [5, 15, 16, 57].

Correlated variability has been shown to be impacted by behavior and brain states. For example, it has been observed that behavior such as running, whisking, and pupil diameter are encoded in V1 and other brain areas [7]. In these contexts, the behavioral subspaces could be estimated directly (as in [7]) and their optimality could be assessed using the FA null model. In experiments with visual attention, it has been shown that attention can modulate both the within-area and between-area correlated variability [31, 58, 59], which can lead to better coding fidelity or better communication of information as assessed by the shuffle null model. Similarly, in an associative task, learning has been shown to modulate the mean response manifold and correlated variability to improve coding in pairs of neurons. The null models developed here could be used to assess whether the modulation due to attention or learning changes the optimality of the correlated variability. Emerging neural recording technologies will allow neuroscientists to simultaneously record from a larger fraction of neurons in a region and more regions, all while the animals are performing naturalistic behaviors. Given these possibilities, the biological origins of correlated variability and how they are modulated by neural circuitry can be further traced and evaluated.

In summary, we find that the geometry of correlated variability in sensory areas leads to highly suboptimal coding for transmission of information about the stimulus. Given the consistency of the findings across datasets, we expect our results would hold true in other organisms, sensory areas, and experimental paradigms. Investigated more broadly, understanding the optimality of correlated variability could lead to a better understanding of the sources of variability is neural circuits and biological constraints that lead to suboptimality. Furthermore, quantitatively evaluating normative theories allows us to adjudicate between competing proposed functions of sensory systems, for example, efficient coding versus predictive information coding.

## Acknowledgements

J.A.L. was supported by the LBNL LDRD “Deep Learning for Science” and the Weill Institute for Neuroscience at UCSF (Bouchard). P.S.S. was supported by by the Department of Defense (DoD) through the National Defense Science & Engineering Graduate (NDSEG) Fellowship Program. M.E.D. was supported by an LBNL LDRD. M.T.S. was supported by the National Science Foundation Graduate Research Fellowship (DGE 1752814). K.E.B. was supported by DOE ASCR (FP00009697), NIH (RNS118648A), the Kavli Institute, the Weill Institute for Neuroscience, and the LBNL LDRD “Coordination of Distributed Cortical Circuits.” This research used resources of the National Energy Research Scientific Computing Center, a DOE Office of Science User Facility supported by the Office of Science of the U.S. Department of Energy under Contract No. DE-AC02-05CH11231. We would like to that the Neural Systems and Data Science Lab and Frederic Theunissen for feedback.

## Methods

### Neural Recordings

We examined correlated variability in a diverse set of datasets, spanning distinct brain regions, animal models, and recording modalities. We used calcium imaging recordings from mouse retinal ganglion cells, single-unit recordings from macaque primary visual cortex, and micro-electrocorticography recordings from rat auditory cortex. We briefly describe the experimental and preprocessing steps for each dataset. See **Figure 1** and **Table 1** for summaries of the datasets.

**Table 1:**
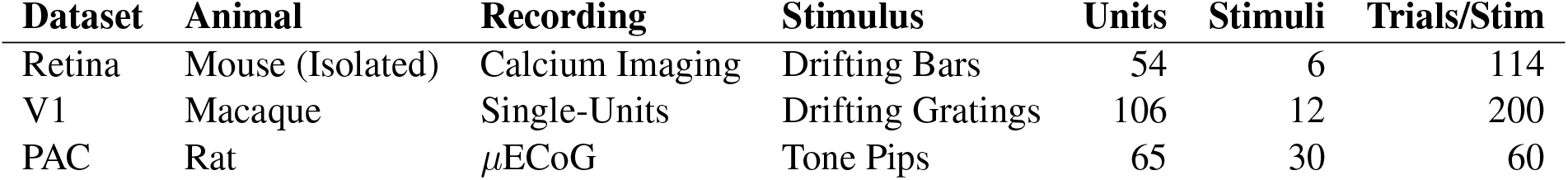
Experimental dataset summary.

### Recordings from mouse retina

Mouse retina data was collected via ex vivo 2-photon calcium imaging in an isolated retina preparation [1]. The retina was bulk loaded with Cal-520 AM dye using a previously described multicell bolus loading technique [2], and then imaged with ScanImage software [3] at 2.96 Hz in the ganglion cell layer of a 425 x 425 *μ*m area of ventral retina. Visual stimuli were delivered via an ultraviolet LED (375 nm) coupled to a digital micromirror device, and were presented on the flyback of the fast-axis scanning mirror during a scan to interleave the stimuli with imaging [1, 4]. Visual responses were elicited via 600 × 600 *μ*m bars drifting for 2.93 s at 750 *μ*m/s in one of 6 directions (spanning 0° to 300°), with a 5 second intertrial interval. Each direction was presented 114 times, for a total of 684 trials per cell. Fluorescence signals from 832 manually selected regions of interest were baseline subtracted and normalized to calculate a ∆F/F_0_ time series. Of these regions of interest, 54 were used for further analysis after determination of directional tuning via permutation testing and manual screening. Per-trial RGC activity used in the analysis here is the maximum ∆F/F_0_ value. Retina data was collected by Summers. Further details on surgical, experimental, and preprocessing steps can be found at [4, 5].

### Recordings from macaque primary visual cortex (V1)

Primary visual cortex data (V1) was comprised of spike-sorted units simultaneously recorded in anesthetized macaque monkey. The data was obtained from the Collaborative Research in Computational Neuroscience (CRCNS) data sharing website [6] and was recorded by Kohn and Smith [7]. This dataset contains recordings from three monkeys, of which the main text presents results from the first one (see Appendix for results on additional two monkeys). Recordings were obtained with a 10 × 10 grid of silicon microelectrodes spaced 400 *μ*m apart and covering an area of 12.96 mm^2^. The monkey was presented with grayscale sinusoidal drifting gratings, each for 1.28 s. Twelve unique drifting angles (spanning 0° to 330°) were each presented 200 times, for a total of 2400 trials per monkey. Spike counts were obtained in a 400 ms bin after stimulus onset. A total of 106 units were isolated in the monkey presented in the main text. These units were chosen by the original authors such that *i*) their signal-to-noise ratio (the ratio of the average waveform amplitude to the standard deviation of the waveform noise) was at least 2.75, *ii*) the best grating stimulus evoked at least 2 spikes/s, and *iii*) the variance-to-mean response ratio did not exceed 10. Further details on the surgical, experimental, and preprocessing steps can be found in [8, 9].

### Recordings from rat primary auditory cortex (PAC)

Auditory cortex data (PAC) was comprised of cortical surface electrical potentials (CSEPs) recorded from rats with a custom fabricated micro-electrocorticography (*μ*ECoG) array. The *μ*ECoG array consisted of an 8 × 16 grid of 40 *μ*m diameter electrodes. Anesthetized rats were presented with 50 ms tone pips of varying amplitude (8 different levels of attenuation, from 0 dB to 70 db) and frequency (30 frequencies equally spaced on a log-scale from 500 Hz to 32 kHz). We only used samples for the lowest 3 levels of attenuation since these evoked the largest responses. Each frequency-amplitude combination was presented 20 times, for a total of 3 × 30 × 20 = 1800 samples. The response for each trial was calculated as the *z*-scored to baseline, high-*γ* band amplitude of the CSEP, calculated using a constant-Q wavelet transform. The maximum of the per-trial high-*γ* activity was used in the analysis here. Of the 128 electrodes, we used 65, selecting those that recorded from primary auditory cortex. Data was recorded by Dougherty & Bouchard. Further details on the surgical, experimental, and preprocessing steps can be found in [10, 11].

### Linear Fisher information measures coding fidelity

A commonly used measure of coding fidelity in the context of decoding is the Fisher information, which provides a limit on how accurately a readout of a neural representation can be used to determine the value of the stimulus [12]. Formally, the Fisher information is a lower bound on the variance of an unbiased estimator for the stimulus. In practice, the Fisher information is analytically intractable. An alternative measure is the linear Fisher information (LFI), defined in Equation 1. The LFI acts as a suitable lower bound to the Fisher information and is the most commonly used measure of coding fidelity in correlated variability analyses [13–20].

Experimental neuroscience datasets only consider discrete sets of stimuli, which are not amenable to the computation of LFI as posed in Equation 1. In particular, the derivative of the average neural activity must be estimated by considering the neighboring pairs of stimuli. Thus, in practice, we calculate the coarsened linear Fisher information [21], which is defined for two stimuli *s*_1_ and *s*_2_ as

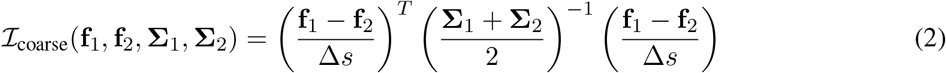

where f_1_ = f (*s*_1_), f_2_ = f (*s*_2_), Σ_1_ = Σ(*s*_1_), Σ_2_ = Σ(*s*_2_), and ∆*s* is the stimulus difference between *s*_1_ and *s*_2_, whose form may depend on the stimulus structure. In addition, we use the unbiased LFI estimator [20] for the observed LFI values as well as for the sampled from null models. Note that since the corrections to the naïve estimator only depend on the dimensionality of the neural population and number of samples, the corrections only impact the raw LFI values and not percentiles. In this work, we use the terms “coarsened LFI” and “LFI” interchangeably.

### Assessing the optimality of neural data with null models

Information theoretic analyses of neural data often ask whether the observed neural data is “optimal.” In the case of correlated variability, the question can be posed as: are the observed covariances optimal from a decoding perspective? Here, we will quantify the coding fidelity with the linear Fisher Information (LFI, Eq. 1)? In this case, LFI can be infinitely large if Σ → 0 (or at least if the subspace of Σ^−1^ defined by 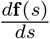 diverges). This answer is likely unsatisfying because neural systems have many sources of variability, and so expecting a neural system to become noiseless or exactly remove noise from a subspace seems implausible. Therefore, when assessing the optimality of correlated variability, one must decide which aspects of the correlated variability the neural system could modify and which aspects will remain fixed.

In this section, we develop the formalism that will allow us to assess the optimality of observed correlated neural variability. The formalism consists of first defining a covariance parameterization for Σ, which is composed of constraints (fixed parameters) and degrees-of-freedom (free parameters). These constraints and degrees-of-freedom define the space of allowed correlated variability. Ideally, these constraints and degrees-of-freedom have some biological interpretation, e.g., fixed private variability or input from other regions of the brain [22, 23]. Then, a null model is defined by combining a covariance parameterization with a null distribution over the degrees-of-freedom. The distribution of some measure, such as the LFI, under the null model can be used to assess the optimality of the observed neural data.

We first review the commonly used fixed-marginal constraint for correlated variability using our formalism then define the commonly used shuffle and novel uniform correlation null models. Finally, we propose the factor analysis covariance parameterizations and associated null model for assessing optimality which has more biological interpretability. In the following sections we will use the following terminology which we define here:

- **Covariance Parameterization:** a parameterization of Σ which can combine various constraints (fixed parameters) and degrees-of-freedom (free parameters).
- **Constraints:** elements of the covariance parameterization which are estimated from data and fixed.
- **Degrees-of-Freedom:** elements of the covariance parameterization which can potentially be modified or optimized to analyze a null model or optimality.
- **Optimality:** values for the degrees-of-freedom in a covariance parameterization which maximize a specified objective. Here we assess optimality using the Linear Fisher Information (LFI), although this formalism can be applied to other objectives.
- **Null Distribution:** distribution of a covariance parameterization’s degrees-of-freedom.
- **Null Model:** combines a covariance parameterization with a baseline or uniform correlation null distribution over the degrees-of-freedom.

The standard constraint considered for understanding correlated neural variability is to keep the per-neuron marginal distributions fixed. Since the LFI only depends on the covariance of the correlated variability, the fix-marginal parameterization is equivalent to constraining the per-neuron variances to be constant (equivalently, the diagonal of **Σ** is kept constant, diag(**Σ**) = ***σ***^2^). The corresponding degrees-of-freedom in this parameterization are the positive-definite pairwise correlation matrix, ***ρ***, specifically the symmetric, off-diagonal entries, *ρ*_*ij*_ for *i* ≠ *j*, which can vary. Under this parameterization, the observed covariance structure can be compared to other proposed distributions of correlations.

When considering the structure that generates **Σ**, it is desirable that the constraints and degrees-of-freedom be biologically interpretable. This can be achieved by considering the equations that define the mean-centered, single-trial response in terms of the degrees-of-freedom being considered. For the fixed-marginals parameterization, the distribution of the single-trial responses: **f**_*t*_(*s*), can be written in terms of a multivariate normal distribution with the mean response: **f** (*s*), and where the covariance is the element-wise product of the constrained marginal standard deviations: ***σσ***^*T*^, and the free correlations: ***ρ***,

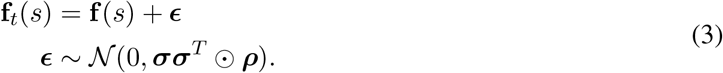

This equation is difficult to directly interpret as a network model, but the correlations could be seen as coming from recurrent activity within the observed neurons.

Given a parameterization (fixed-marginal) and a measure of coding fidelity (LFI), it is possible to find optimal covariance structures as a function of the free parameters. In general, the value (or distribution of values) for the degrees-of-freedom that lead to optimality can be derived analytically or optimized numeri-676 lly. For the fixed-marginal parameterization, this corresponds to finding the points, 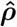, such that

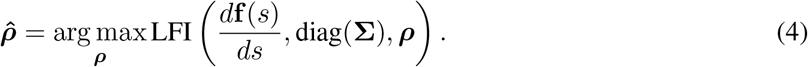

Hu *et al.* [24] characterize the optima of the fixed-marginal parameterization, although they do not provide a constructive way of finding the global optima. We optimize ***ρ*** numerically to find optima. We find that the optimization process finds many local maxima for 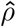 in practice.

### Novel null models allow the assessment of optimality in neural data

So far, we have have laid out a formalism to define the optimal degrees-of-freedom for a specified covariance parameterization. However, it is unlikely that observed neural data will precisely match the predicted optimal degrees-of-freedom, even if the biological system is behaving optimally, so the predictions from Eq 4 cannot be used directly to assess optimality in data. In order to asses the optimality of a observed population of neurons, a null model must be constructed for a corresponding parameterization. In this formalism, constructing a null model corresponds to assuming a null distribution for the degrees-of-freedom of the covariance parameterization. The null distribution should correspond to some notion of “uniform” or “baseline” for the degrees-of-freedom.

For example, the shuffle null model, based on the fixed-marginal parameterization, posits that the baseline distribution of correlations is zero correlations. The shuffle null model compares the LFI of the observed response to the distribution of LFIs where the individual neural responses are independently trial shuffled, that is, with fixed-marginal variability, no underlying pairwise correlations, and empirical pairwise correlations only arising from finite sampling effects. Under this choice of null model, the observed LFI can be beneficial if it has a high percentile under the null distribution which has no correlations. The shuffle null model provides a limited baseline comparison for the observed LFI. In order to assess optimality, the distribution of parameters should be uniform over the space of allowed covariance matrices, which is the motivation for the uniform correlation null model.

Across a population, the median observed percentile across dim-stims can be used to categorize a dataset as optimal: median percentile greater than or equal to 2/3, near-chance: median percentile between 1/3 and 2/3, or suboptimal: median percentile less than 1/3. This categorization is motivated by simplicity in having few categories. However, it is also desirable to not have the optimal and suboptimal categories share a boundary. If they do, small changes in percentiles can switch between optimal and suboptimal. In our case, since the null model defines “near-chance”, having 3 categories is natural. The near-chance boundaries could be set in a number of ways besides the choice for an even division into thirds. A Kolmogorov–Smirnov test could compare the distribution of percentiles to a uniform distribution. However, given the large number of dim-stims we use, empirically, no distributions of percentiles in these datasets would be near-chance for p-value thresholds in sensible ranges. Said another way, almost no empirical distributions of percentiles are statistically similar to a uniform distribution (see **Supplementary Fig. 5a-i** for some example distributions). A looser test could be to test whether a binomial distribution with *p* = 0.5 would lead to the observed distribution of percentiles categorically above and below 0.5. We find that with p-values in sensible ranges this gives comparable boundaries to the division into thirds, but the boundaries differ across datasets due to the variation in the number of dim-stims.

In some cases, it may also be possible to define a distribution over optimal covariances and categorize whether the observed LFI is likely under the optimal covariance distribution. For instance, if there is a unique optimal covariance, the Wishart distribution could be used to create a sampling distribution of optimal LFIs which the observed LFIs could be compared against. This is not possible in our case since there is not generally a unique optimal covariance. This also suffers from the fragility problem by having a boundary directly between optimal and suboptimal.

### Uniform correlation null model

Our first contribution is the uniform correlation null model based on the fixed-marginal parameterization, where the correlations are chosen randomly from a uniform distribution over correlation matrices [25]. This tests whether the observed correlation are optimal with respect to all possible correlations, rather than only comparing against zero correlations. To our knowledge, this null model has not been considered before. Evaluating data under this null model provides a stronger assessment of the optimality of the observed correlated variability than the shuffle null model.

At another extreme, we could attribute all trial-to-trial variability to external sources that the network can shape or filter. To prevent trivial solutions, we can restrict the network to only changing the loading of the variability onto the neurons (through a rotation, **R**). This model was previously discussed [24], but not analyzed due it its incompatibility with the fixed-marginal constraint.

### Factor analysis null model

As a parsimonious combination of the fixed-marginal constraint and pure rotation degrees-of-freedom, we propose using a factor analysis (FA) model to parameterize the correlated variability. Factor analysis decomposes the observed correlated variability into two components: the first is per-neuron private variability, represented as a diagonal matrix 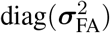, and the second is a low-rank shared variability component, 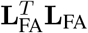, where 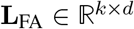, *k* < *d*. We propose that the FA model has private variability and the spectrum of the shared component as constraints and the rotation of the shared components as the degrees-of-freedom, combining aspects of the fixed-marginal and rotation null models. The single-trial response can be written as a function of the mean response: **f** (*s*), private variances: 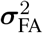, low-rank external sources: **z**_FA_, loading matrix: **L**_FA_, and rotation matrix: **R**

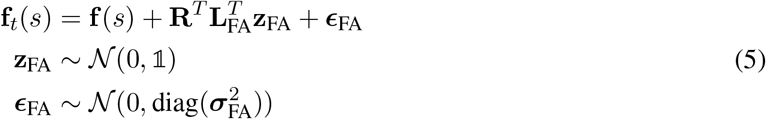

To our knowledge, there is no closed-form solution for 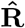 in the FA model to maximize LFI. Instead, to optimize the FA model, the rotation can be numerically optimized by gradient ascent. To construct the FA null model, a uniform distribution (Haar distribution) over special orthogonal rotations [26] is applied to the rotations.

To estimate the initial 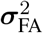 and **L**_FA_, we fit a factor analysis model to the samples [27]. In fitting the model we had two requirements. The first is that we wanted the dimensionality of the shared component, *k* to be as large as possible so that the observed covariance can be modeled as accurately as possible. In opposition to this, we wanted the factor analysis model parameters to be identifiable, meaning the private variance estimate is unique, which places a limit, which depends on *d*, on how large *k* can be [28]. In practice, we find the largest *k* which is lower than the identifiability bound where different initializations return the same parameters. Note that factor analysis is never identifiable in 2 dimensions, so we do not consider *d* = 2.

### Population statistics across dim-stims measure optimality under a null model

Each dataset can be described by a *D × N* design matrix **X**, where *D* is the total number of samples and *N* is the number of units in the population (**Fig. 2f**). We considered distributions of LFI across dim-stims, or sub-components of the design matrix. To create dim-stims, we first selected a dimlet of size *d* by subsampling *d* units from the population at random, resulting in the *D* × *d* design matrix **X**^*d*^ (**Fig. 2f**). Next, we created the dim-stim by further subsampling the design matrix according to a specific stimulus pairing. Specifically, we chose two neighboring stimuli, *s*_1_ and *s*_2_ (**Fig. 2f**), and isolated the samples of **X**^*d*^ corresponding to those stimuli, thereby creating a pair of design matrices 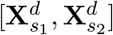. The dim-stim maps to the task of discriminating between two neighboring stimuli using a sub-population’s responses across trials to those stimuli, which can be visualized in the neural space (**Fig. 2g**).

For each dataset, we considered dimlet dimensions *d* = 3 – 20. As we only allowed neighboring stimulus pairings, the number of available stimulus pairings for a dimlet was 6 (retinal), 12 (V1) and 29 (PAC). Note that the retinal and V1 stimulus sets are circular, providing an additional stimulus pairing. In the retinal and V1 datasets, we drew 1,000 dimlets for each dimension *d*, and considered all stimulus pairings per dimlet, resulting in 1, 000 × 6 = 6, 000 dim-stims for the retinal dataset and 1,000 × 12 = 12000 dim-stims for the V1 dataset. To manage computation time, we considered 3, 000 unique dim-stims for the PAC dataset, selecting both the dimlet and stimulus pairing at random for each dim-stim.

For each dim-stim, we calculate its observed LFI, defined as 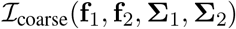. Specifically, we computed

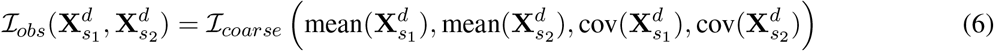

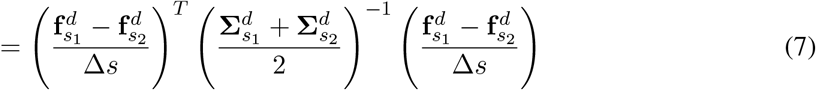

where 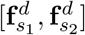 are the dim-stim average responses, 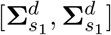 are the dim-stim covariances, and Δ*s* is the stimulus difference, or Δ*s* = |*s*_1_ – *s*_2_|. When necessary, the stimulus difference was taken as a circular difference (retinal and V1 datasets). Since the LFI is scaled by the units of the stimulus difference, it is only meaningful to compare observed LFIs within a particular stimulus type. In this work, since all datasets use a different stimulus the LFIs may not have a meaningful relationship across datasets.

Each null model acts on the design matrices of a dim-stim and outputs a distribution of covariance matrices. For example, the fixed-marginal null model shuffles the data within the design matrix, producing new design matrices 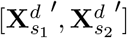 and corresponding covariances 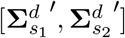. We then calculate the LFI using the new covariance matrices. Each null model can be summarized as such: a sampled transformation is applied to the observed dim-stim, producing new sampled covariance matrices and therefore a sample of LFI from the null. The shuffle null model transformed the data directly, so we write its LFI as

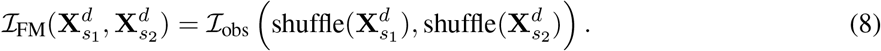

Meanwhile, the uniform and factor analysis null models transform the covariance parameterization directly, so we write their LFIs as:

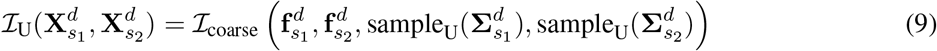

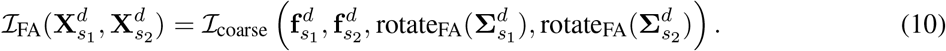

Equations 8 and 10 capture a single application of a null model. Specifically, shuffle(·) shuffles the neural data, sample_U_(·) samples a random off-diagonal correlation structure and applies it to the covariance, and rotate_FA_(·) applies a rotation to the shared component of the covariance. However, we were interested in characterizing the entire distribution of the null model. Thus, for each dim-stim, we applied 1, 000 samples of the null model to obtain a null model distribution of LFIs. We then calculated observed percentiles as the fraction of samples for which the observed LFI exceeded the null model LFI. Thus, each observed dim-stim has its own corresponding observed percentile, per null model.

When summary statistics are reported such as the median LFI, median percentile, or the optimal fraction, 95% bootstrap confidence intervals from 1,000 bootstrap resamples are reported [29].

### Optimal fraction calculation

The optimal fraction of a population was calculated in the following way. Given a set of dim-stims at a particular dimlet dimension, the observed percentiles were calculated for each dim-stim. Then, the percentiles were sorted from largest to smallest. The optimal fraction of the percentiles is initialized as the largest single percentile. Starting from this initialization, the median percentile of the current optimal fraction is calculated. If the median is greater than or equal to 2/3, the next smallest percentile is included in the optimal fraction and the process continues to iterate. If the optimal fraction is less than 2/3, the process terminates. This defines the largest possible fraction of the percentiles that can be retained and have their median be greater than or equal to 2/3. For reference, the top 2/3 of a uniform distribution (i.e., [1/3, 1]) of percentiles has median equal to 2/3.

### Measures of biological plausibility

We calculated the mean Fano factors (FF) for a dim-stim, based on the per-unit variance and response means

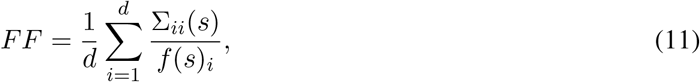

of the observed and optimal covariances matrices directly from the mean response and covariance matrix parameters (Supplemental **Fig. 2**).

We calculated the negative density (ND) as follows. For each dim-stim, we calculated 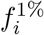, the neural activity at the 1st percentile, for each neuron *i*. We then computed 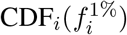, the cumulative density at 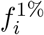 for a Gaussian obtained from either the observed covariance or the optimal covariance under the null model (Supplemental **Fig. 2**, shaded regions in marginals). The ND, then, was defined as the maximum CDF_*i*_ among the neurons in the dimlet (Supplemental **Fig. 2**, dark gray shaded regions).

### Distance and tuning ranking dim-stims for subselection

For the retina and PAC datasets, we have access to the spatial locations of the RGC/electrode. For distance-based subselection, we compute the average pairwise distance between neural units for each dim-stim. The dim-stims are ranked by this distance and the 10% of dim-stims with the smallest average distance are subselected.

For tuning-based subselection, the stimuli are ranked for each neural unit based on the mean neural activity (tuning). The rank was used because is less sensitive to absolute firing rates compared to using the activity per stimuli, which would biased the subselection towards dim-stims which contain neural units with high firing rates. We then sort the dim-stims by their average tuning rank across dimlets and calculate percentile statistics for the 10% of dim-stims that have the highest tuning ranking.

## Appendix Geometric contributions to neural correlated variability

The geometry of three types of potential contributions to neural variability are shown. Private variability is a zero-correlation contribution (Fig. 1a) [1]. Shared variability can be a low-rank contribution whose orientation depends on the synaptic loading onto the observed neural units (Fig. 1b) [2, 3]. Differential correlations lie along the 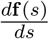 direction (Fig. 1c) [4].

**Figure 1:**
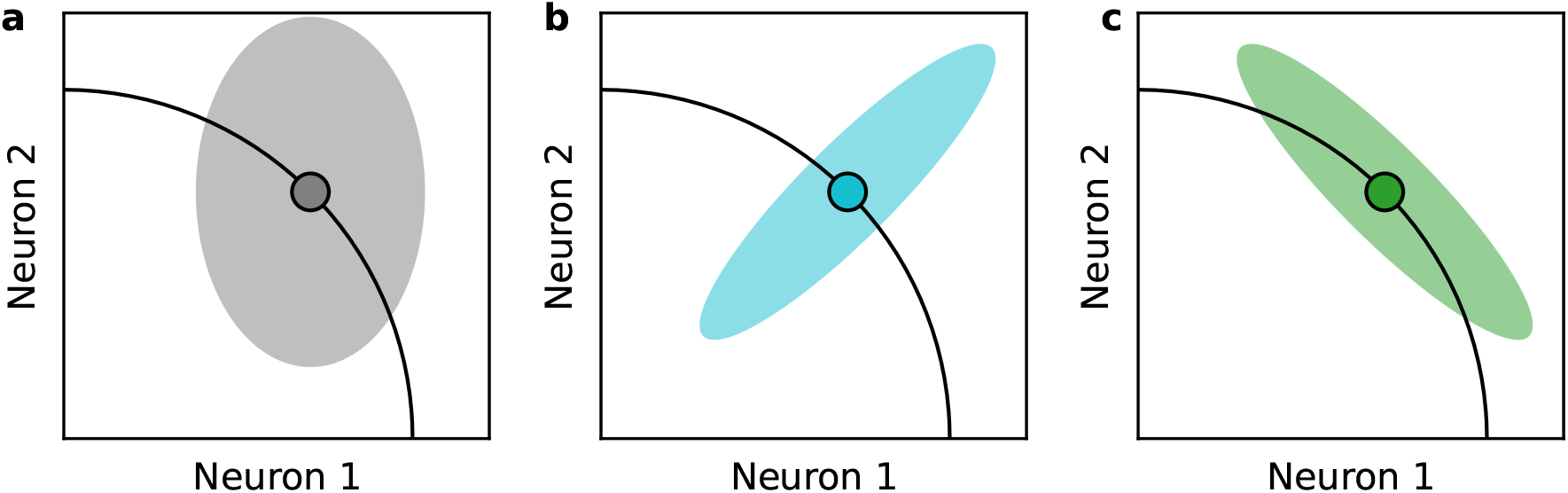
Geometric contributions to neural correlated variability. Each plot depicts the neural space, whose axes correspond to the activities of a specific pair of neurons to a stimulus. Black curves denote the mean responses across different stimuli (i.e., tuning curves). Variability about a specific stimulus mean activity (solid points) may exhibit: **a.** Private, uncorrelated variability in each neural dimension, **b.** Correlated variability, with correlations in the neural space, and **c.** Differential correlations, which lie parallel to the mean activity curve.

## Measures for assessing biological accessibility

Consider an example V1 dim-stim for a dimlet of size *d* = 3, with low observed percentiles under both the null models (e.g., *p*_U_ = 0.001 and *p*_FA_ = 0.0). We plot the observed covariance structure, projected into two neural dimensions, in Figure 2a (black covariance denotes average covariance). Next, we compare the observed structure to that of the optimal structure, both within the factor analysis null model (Fig. 2b) and the uniform null model (Fig. 2c).

**Figure 2:**
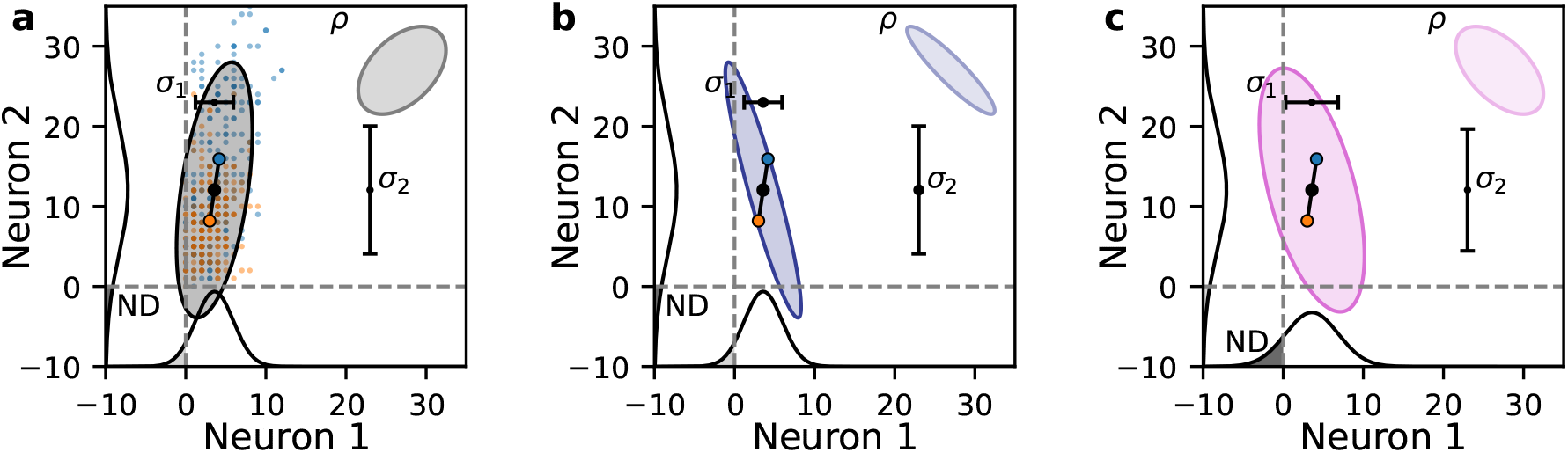
Measures for assessing biological accessibility. Data, fit and optimal covariances (at 2 standard deviations) are from a *d* = 3 dimlet-stim projected into the first 2 neurons. The marginal probabilities of the multivariate Gaussian fits are shown along the axes and the areas with values less than the empirical 1% are shaded grey with the maximum excess negative density in dark grey (annotated with “ND”). The marginal means and standard deviations (for Fano factor calculations) are shown in the black error bars (annotated with “*σ*” and neuron number). For each covariance, the corresponding correlation ellipse (*ρ*, with an arbitrary uniform scaling) is shown in the top right of the plot. **a:** Observed single-trial neuron responses to stimuli 1 and 2 (orange and blue dots) and the respective means (outlined circles). Their joint meant is the black circle and the observed mean covariance is in gray. **b:** Covariance and marginals from an optimal fixed-marginal correlation. **c:** Covariance and marginals from the optimal Factor Analysis rotation.

The observed correlated variability structure (Fig. 2a) exhibits poor discriminability, because a large amount of variability is oriented parallel to the stimulus manifold (Fig. 2, black lines in the empirical covariance ellipse). We consider several measures of biological plausibility for the optimal covariances. The first is the median absolute correlation of the optimal covariances (Fig. 2, ellipse labeled *ρ* in top right shows optimal correlation), which is most relevant for the uniform correlation null model. The second is is the Fano factors (FF) of the optimal covariance relative to the Fano factors of the observed covariance (Fig. 2, black mean and standard deviation indicators labeled with *σ*_1_ and *σ*_2_). The third is the cumulative marginal probability the optimal covariance has below the 1st percentile of the observed data (Fig. 2, gray regions in marginal distributions labeled ND, negative density). These measures only take on a limited range of values in measured neural activity, and may impede a neural system from obtaining an optimal correlated variability structure. The uniform correlation null model preserves the per-RGC/neuron/electrode mean and variance, and so the FF and ND measures are only relevant for the factor analysis null model.

However, the optimal covariance orientations for the factor analysis model may possess different Fano factors (Fig. 2c). Thus, we aimed to assess whether biologically unachievable Fano factors shared any relation with the sub-optimality exhibited by the neural codes in our analyses. We summarized each dim-stim with an aggregate Fano factor, by averaging the Fano factors of that dim-stim’s individual units. We repeated this process for the optimal noise covariances under each null model, using the variances from the diagonal of the optimal noise covariance matrix directly when calculating Fano factors.

To quantify this phenomenon, we calculated the absolute difference in negative density (ND), which captures the degree to which an optimal covariance puts differing cumulative density in the negative response space (typically higher density). Thus, a larger ND implies that the covariance places an excess of density in the negative or low-activity regions for at least one dimension of the neural space. On the other hand, a lower ND is more biologically plausible, as this implies there is less negative density, although Gaussian fits will always put some non-zero density in the negative.

## Optimal correlations for the fixed-marginal parameterization lie on the boundary of possible correlations

Hu *et al.* [5] show analytically that optimal covariances in the fixed-marginal parameterization lie on the boundary of allowed correlations, which will generally have large absolute pairwise correlations. We reproduce this results computationally. For each dim-stim, we compare the 90% percentile of the off-diagonal entries absolute correlation matrix for the observed covariance matrix, the optimal uniform correlation (UC) null model matrix, and the optimal factor analysis (FA) null model matrix. The histograms across dim-stims for 4 dimensions is shown in Figure 3. The observed 90% abs. correlations are rarely larger than 0.7. The FA optimal 90% abs. correlations have a larger spread towards higher correlations, but do not have density at 1. However, the UC optimal covariance have 90% abs. correlations that consistently have peaks in probability mass at 1.

**Figure 3:**
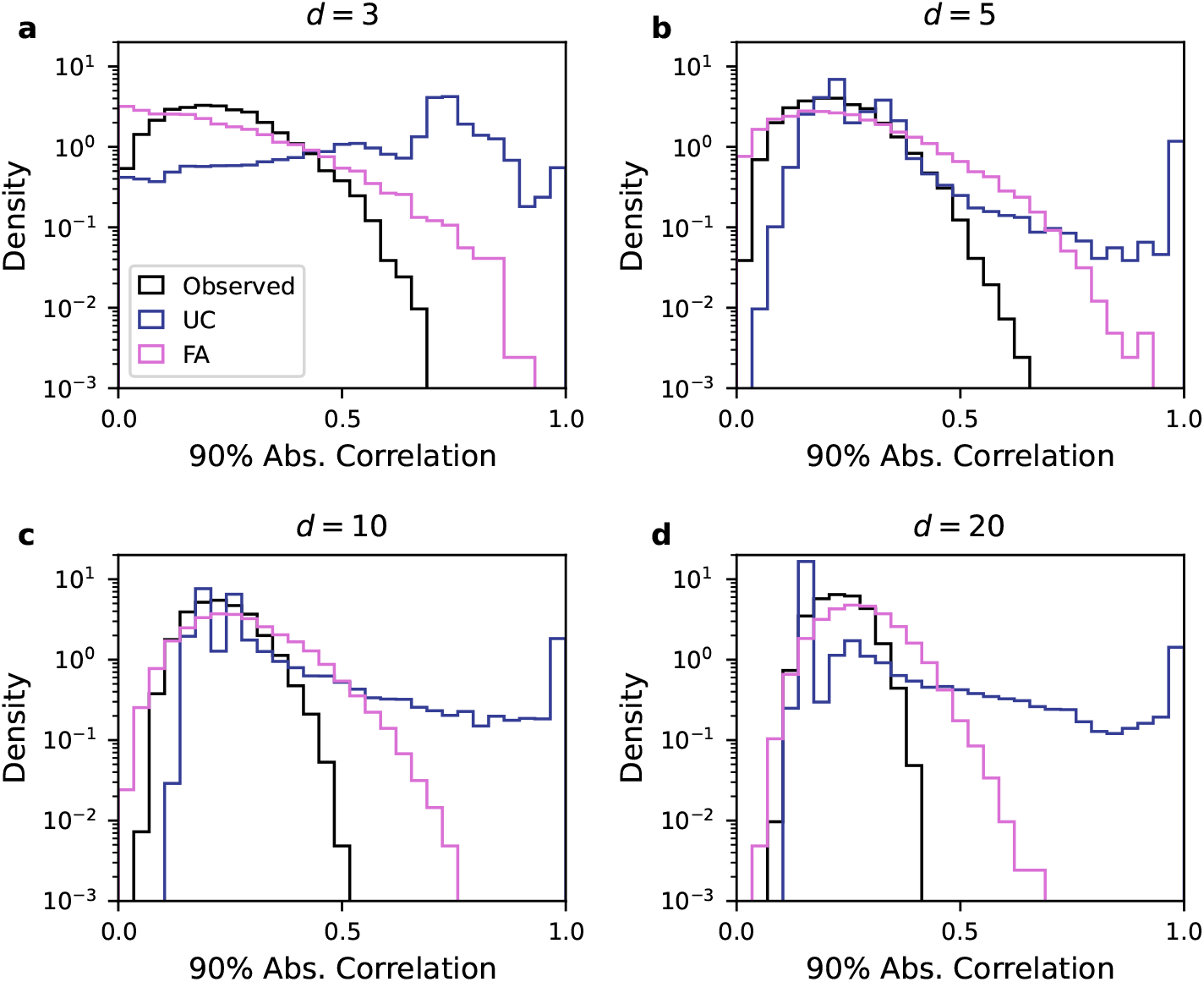
Optimal correlations for the fixed-marginal parameterization lie on the boundary of possible correlations. For each dimension, *d*, the 90th percentile of the absolute value of the pairwise correlations is histogrammed across dim-stims. Color indicates whether the statistic is from the observed covariance or optimal null model covariance. **a-d.** Dimensions 3, 5, 10, and 20, respectively are shown.

## Biologically motivated subselection of dim-stims remains suboptimal

In the main text, biological subselection in Figure 5 was done based on the distance- and tuning-based criteria as they might correspond to biological criteria enforced during development or learning. It is also possible to subselect the dim-stims using the Fano factor (FF) and negative density (ND) criteria directly for the factor analysis null model. Here, we compute the average rank the dim-stims based on their violation of the FF and ND criteria and retain the 10% of dim-stims with the least average violation. This criteria leaves the population percentiles suboptimal (Fig. 4a-c).

**Figure 4:**
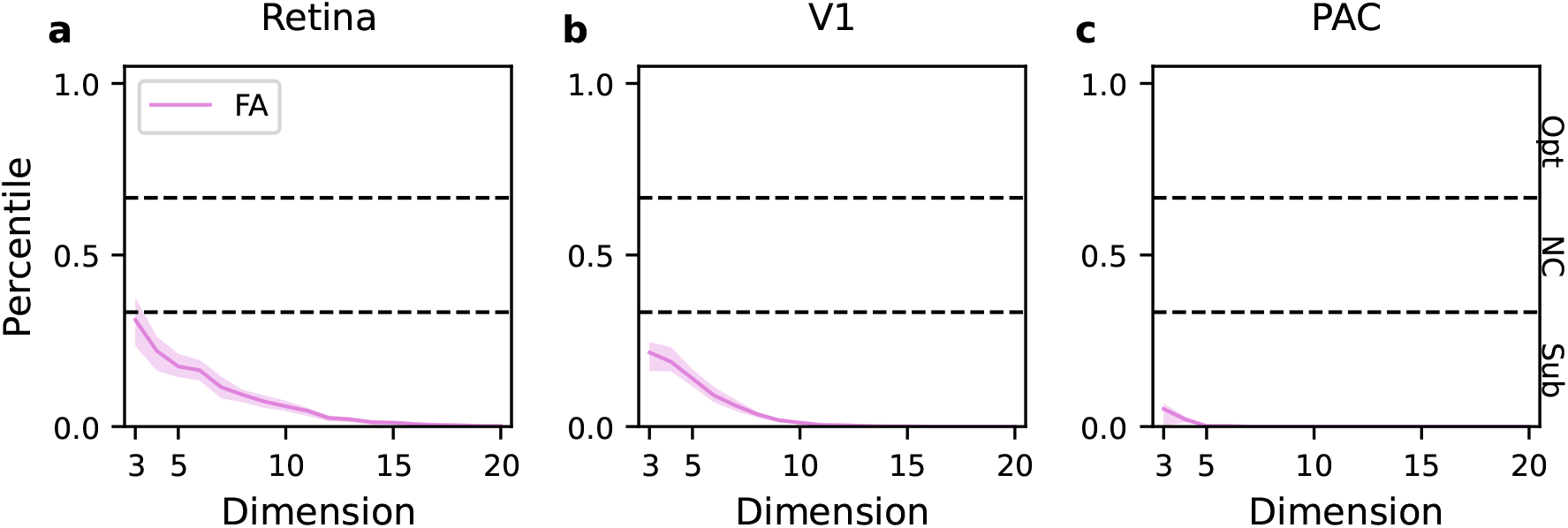
Biologically motivated subselection of dim-stims remains suboptimal. The dim-stims are subselected based on biological criteria and their median percentiles are shown as a function of dimension. Black dashed lines indicate the 33rd-66th percentile range. Shaded regions bound the 40th to 60th percentiles of the subselected percentile distributions. **a-c.** For the FA null model, dim-stims were subselected to minimize their average Fano factor and ND deviations (0th-10th percentile). The median and 33-66% of the percentiles for this subpopulation is shown.

## There is an exponentially small peak of optimal dim-stims for the factor analysis null model

For the factor analysis null model, there is sometimes a peak of percentiles near 1 (Fig. 5a-i). For some dimensions, the peak has higher density than what would be expected from a uniform distribution. To calculate the peak width at each dimension, the percentiles are sorted and, starting from the largest percentiles, the observed percentiles are compares with the percentiles that would be expected from a uniform distribution. The peak width is the fraction of percentiles corresponding to the smallest percentile that has a value larger than what is expected from a uniform distribution. Across datasets, the peak width is exponentially small as a function of neural dimension (Fig. 5j-l). The uniform correlation null model does not have peaks near 1 for any dimension or dataset.

**Figure 5:**
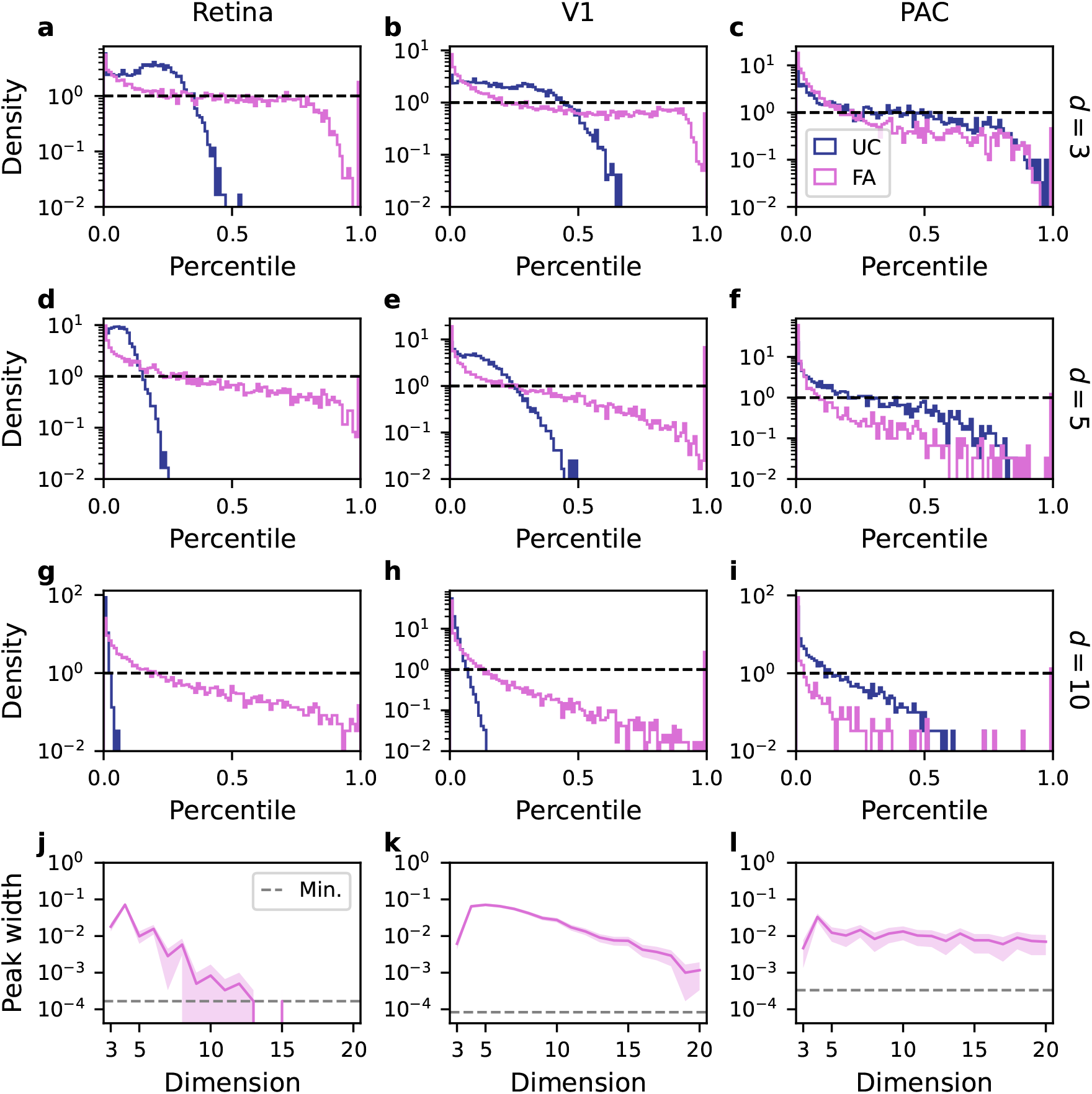
There is an exponentially small peak of optimal dim-stims for the factor analysis null model. **a-i.** The histograms of the percentiles distributions are shown across datasets and null models for dimensions 3, 5, and 10 (in rows). Black dashed lines indicate the density of a uniform distribution. Note the y-axis is log-scaled. **j-l.** Across dimensions, the width of the greater-than-uniform peak is shown. Shaded regions are the 95% CI for the peak widths. Gray dashed line indicates the minimum non-zero peak width that can be estimated due to finite sampling.

## V1 datasets give similar results across monkeys

The PVC11 dataset (here V1) from CRCNS has data from 3 different monkeys [6]. In the main text, we used monkey 1. Although there are differences in the distribution of pairwise correlations (Fig. 6a), they do not lead to qualitative differences in the results from the main text across animals. Figure 6b-m reproduce the main results for all 3 monkeys.

**Figure 6:**
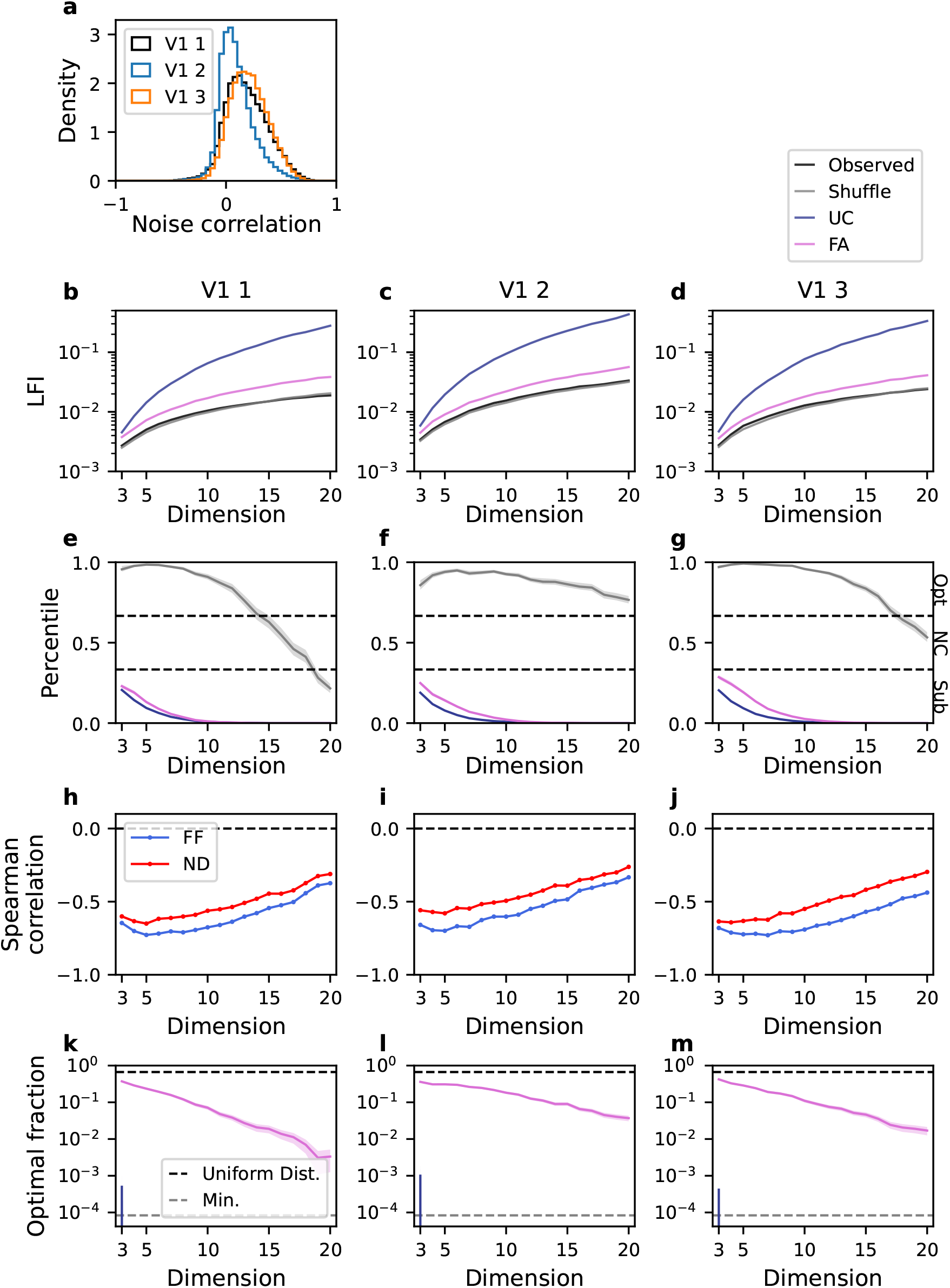
V1 datasets give similar results across monkeys. Main text results are repeated for monkeys 2 and 3 (V1 2 and V1 3, second and third columns) and compared with monkey 1 (V1 1, first column) which is reproduced here. Panel **a** corresponds to main text Figure 1. Panels **b-g** correspond to main text Figure 3. Panels **h-j** correspond to main text Figure 4. Panels **k-m** correspond to main text Figure 5. See main text for panel details.

